# Discovery of a new activator of Slack potassium channels with robust efficacy in models of histamine-independent and chronic itch

**DOI:** 10.1101/2023.10.05.560997

**Authors:** Annika Balzulat, W. Felix Zhu, Cathrin Flauaus, Victor Hernandez-Olmos, Jan Heering, Sunesh Sethumadhavan, Mariam Dubiel, Annika Frank, Amelie Menge, Maureen Hebchen, Katharina Metzner, Ruirui Lu, Robert Lukowski, Peter Ruth, Stefan Knapp, Susanne Müller, Dieter Steinhilber, Inga Hänelt, Holger Stark, Ewgenij Proschak, Achim Schmidtko

## Abstract

Various disorders are accompanied by histamine-independent itching, which is often resistant to the currently available therapies. In this study, we hypothesized that pharmacological activation of Slack (Kcnt1, K_Na_1.1), a potassium channel highly expressed in itch-sensitive sensory neurons, has therapeutic potential for the treatment of itching. Based on the Slack-activating antipsychotic drug, loxapine, we designed a series of new derivatives with improved pharmacodynamic and pharmacokinetic profiles that enabled us to validate Slack as a pharmacological target *in vivo*. One of these new Slack activators, compound 6, exhibited negligible dopamine D_2_ and D_3_ receptor binding, unlike loxapine. We found that compound 6 displayed potent on-target antipruritic activity in multiple mouse models of acute histamine-independent and chronic itch without motor side effects. These properties make compound 6 a lead molecule for the development of new antipruritic therapies targeting Slack.

## 1. Introduction

Itching (also known as pruritus) is defined as an unpleasant sensation that evokes a desire to scratch. Depending on whether histamine release is involved, itching can be broadly divided into histamine-dependent (histaminergic) and histamine-independent (non-histaminergic) itching.^[1–3]^ Like pain, acute itching serves as an important protective mechanism for detecting potentially harmful stimuli. However, chronic itch (i.e., persisting for more than 6 weeks in humans) no longer serves a useful function but instead imposes suffering and may compromise the quality of life to a degree often comparable to that seen in chronic pain.^[4]^ Chronic itch is a debilitating symptom that accompanies various skin disorders, systemic diseases, such as chronic kidney or cholestatic liver disease, psychiatric diseases, and neuropathies, or is idiopathic. An estimated one third of dermatology patients and nearly 15% of the general population experience chronic itching.^[5–7]^ As most types of chronic itch are histamine-independent and, therefore, resistant to antihistamines, there is an urgent need to develop novel treatment strategies.^[8]^

Most known mechanisms of pruriception (itching sensation) begin with the activation of itch-sensitive sensory neurons. Several key receptors, including members of the Mas-related G-protein-coupled receptor (Mrgpr) families, are important for detecting non-histaminergic chemical itch signals.^[2, 9]^ Recent single-cell RNA-sequencing (scRNA-seq) studies have revealed that itch-related genes are highly expressed in distinct subpopulations of sensory neurons. For example, in a pioneering scRNA-seq study by Usoskin et al.,^[10]^ 11 principal types of sensory neurons were identified, of which three non-peptidergic populations (NP1, NP2, and NP3) were proposed to be itch-sensitive. After activation, these sensory neurons signal to the dorsal horn of the spinal cord, where ongoing information is further processed and transmitted to itch-signaling pathways ascending to the brain.^[11]^

The excitability of sensory neurons is driven by various types of ion channels, among which K^+^ channels are the most diverse class governed by more than 75 genes in humans. Owing to their diversity and tissue-dependent expression patterns, they are increasingly recognized as potential drug targets.^[12, 13]^ Notably, the K^+^ channel Slack (also known as K_Na_1.1 or Slo2.2; gene *Kcnt1*), which is activated by intracellular Na^+^ and inhibited by bivalent cations,^[14–16]^ is highly enriched in the itch-sensitive NP1, NP2, and NP3 subsets of sensory neurons of the peripheral nervous system in mice^[10, 17]^ (Figure S1A,B, Supporting Information), suggesting that Slack has a role in itch. In fact, a remarkable functional role of Slack in pruriception is reflected by the increased scratching behavior of Slack knockout mice after exposure to pruritogens such as chloroquine.^[18]^ These findings in combination with the observation that Slack expression in non-human primate sensory neurons (Figure S1C, Supporting Information) and human sensory neurons (Figure S1D, Supporting Information) is enriched in itch-associated cell populations^[19, 20]^ support the hypothesis that pharmacological activators of Slack hold therapeutic potential for the treatment of itch.

A previous study with a library screen of pharmacologically active compounds reported that the first-generation antipsychotic drug loxapine activates Slack.^[21]^ Interestingly, in preliminary experiments, we observed that systemic administration of a low dose of loxapine considerably alleviated chloroquine-induced scratching behavior in wildtype mice but did not affect scratching behavior in Slack knockout mice, indicating that the loxapine-induced antipruritic effect depends on Slack activation. However, the clinical use of loxapine is limited by the typical adverse events associated with first-generation antipsychotic drugs, which are mainly caused by the blocking of dopamine receptors and other neurotransmitter receptors.^[22, 23]^ Hence, in this study, we aimed to develop novel Slack-activating compounds with an improved pharmacological profile as compared to that of loxapine, in particular by reducing off-target activity towards dopamine receptors. We hypothesized that these aims could be achieved by introducing modifications to the loxapine scaffold. Using these new compounds, we aimed to validate Slack as a novel target for itch treatment.

## 2. Results

### 2.1. Development of Slack-activating compounds

We developed a series of 114 new derivatives of loxapine. The new compounds were synthesized using linear synthesis routes (Figure S2, Supporting Information). Their structural features are listed in Table 1. The loxapine scaffold contains a tricyclic core and a piperazine ring substituted with an alkyl residue. Starting from methyl esters of salicylic acid and 2-fluoronitrobenzene derivatives, diarylethers were obtained via nucleophilic aromatic substitution. Subsequent reduction of the nitro group by tin(II)chloride enabled intramolecular amide bond formation. Alternatively, an amide bond was first formed between 2-fluorobenzoic acids and 2-aminophenol derivatives, followed by intramolecular nucleophilic aromatic substitution to afford the same lactam core. The lactam was then converted to the imidoyl chloride using POCl3. In the final step, the substituted piperazines displaced the chloride to afford the desired loxapine derivatives.

**Table 1.**
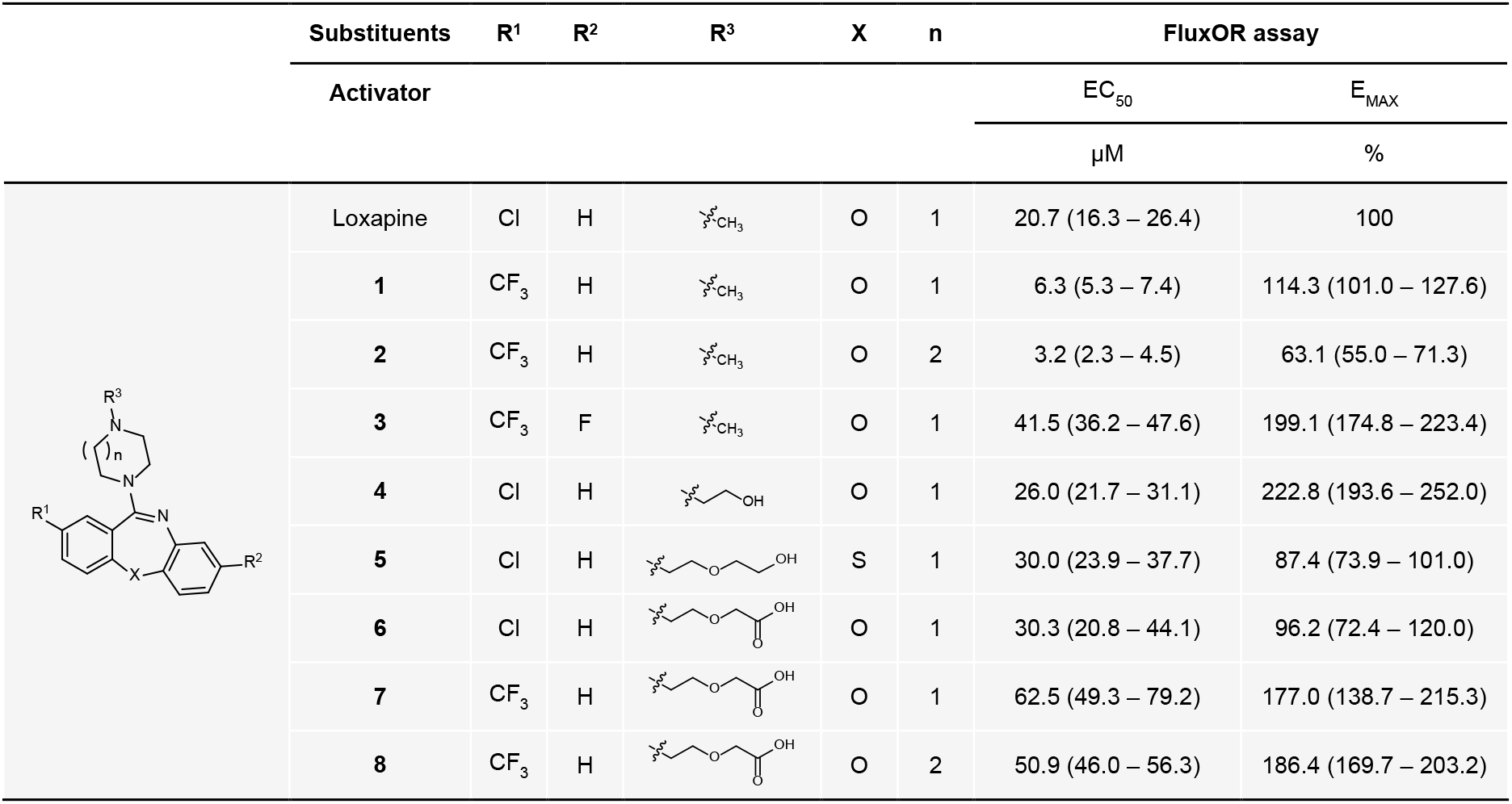
Structural features of the most promising new compounds and their Slack-activating properties. The chemical synthesis scheme of the new compounds is depicted in Figure S2 (Supporting Information). Potency (EC_50_) and efficacy (E_MAX_; % relative to loxapine) values are from a FluxOR assay in HEK-Slack cells, shown in Figure 1A. New compounds were measured in triplicate. EC_50_ and E_MAX_ values are presented as mean with 95% confidence interval.

The functional activity of this series of compounds against human Slack was determined using a cell-based FluxOR potassium ion channel assay in cultured HEK293 cells stably expressing human Slack (herein referred to as HEK-Slack cells) and a Na^+^-free buffer to prevent compound-independent Slack activation by monovalent cations.^[14, 24]^ Having established that the FluxOR assay is capable to detect Slack activation by loxapine (Figure S3, Supporting Information), we conducted concentration-response experiments to determine the EC_50_ and E_MAX_ values. Of the 114 newly synthesized compounds, 31 acted as Slack activators to varying extents. Accordingly, we deemed eight of these as the most interesting candidates. Their Slack-activating properties are listed in Table 1. Potency (EC_50_) values ranged between 3.2 and 62.5 µM, whereas loxapine activated Slack with a potency of 20.7 µM (Table 1 and Figure 1A). Efficacy (E_MAX_) values ranged from 63.1% to 222.8% normalized to Slack activation by loxapine (Table 1 and Figure 1A). Considering that low doses of loxapine exerted Slack-dependent effects in mice,^[24]^ we concluded that the potency of these eight new compounds may be sufficient to activate Slack at standard doses *in vivo*.

**Figure 1.**
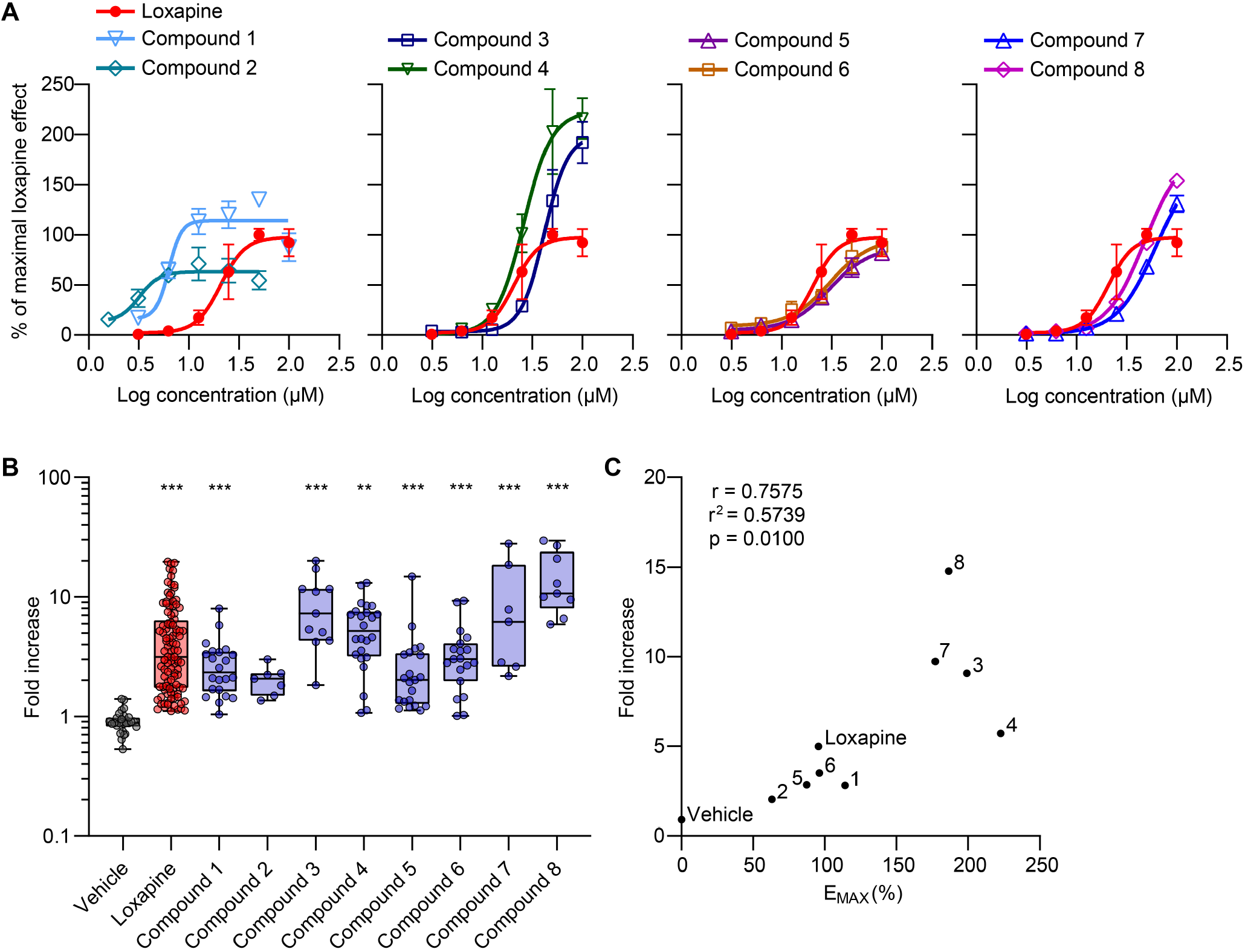
Compounds serving as activators of Slack channels. A) Concentration-response experiments with our new compounds in the FluxOR potassium ion channel assay. HEK293 cells stably expressing human Slack (HEK-Slack cells) were assayed for a potassium channel-mediated thallium response using FluxOR. Each compound was incubated at six concentrations. For each sample, the fluorescence value was calculated and then normalized to the maximum fluorescence value of loxapine. For better comparability, data from the loxapine measurements are presented in all four graphs. Each data point is the average of 3 replicates. Data represent the mean ± SD. Resulting EC_50_ and E_MAX_ values from these measurements are shown in Table 1. B) Patch-clamp recordings confirm that new compounds evoke Slack-mediated potassium currents. Whole-cell voltage recordings on HEK-Slack cells were performed at baseline and after incubation with a new compound (50 µM), loxapine (50 µM) or vehicle (external solution containing 0.03% DMSO). Loxapine was incubated in each series of experiments as a positive control. Shown is the fold increase in current densities (pA/pF) elicited by new compounds, loxapine, and vehicle relative to baseline at a voltage of +80 mV (vehicle, *n* = 28 cells; loxapine, *n* = 104; compound 1, *n* = 22; compound 2, *n* = 7; compound 3, *n* = 10; compound 4, *n* = 24; compound 5, *n* = 21; compound 6, *n* = 19; compound 7, *n* = 7; compound 8, *n* = 9 cells; \*\**P* < 0.01, \*\*\**P* < 0.001 vs. vehicle; Kruskal-Wallis test with Dunn’s correction. Box-and-whisker plots represent maximum and minimum values, and the box shows the first, second (median) and third quartile values. Current-voltage curves from these recordings are presented in Figure S4 (Supporting Information). C) Correlation of relative E_MAX_ values of new compounds obtained in the FluxOR assay (presented in A and Table 1) with relative current densities obtained in the patch-clamp experiments (presented in B). Statistical significance was assessed by a Pearson correlation.

The evaluation of analogs in our series of Slack activators in the FluxOR assay suggested that replacement of the chloro substituent at R^1^ by a trifluoromethyl group improved activity (compound 1). Additional introduction of a fluoro substituent at the other aryl ring (R^2^) increased efficacy but reduced potency (compound 3). In some cases, the sulfur analogs showed comparable activity (compound 5). Replacement of piperazine with homopiperazine improved the EC_50_ value but decreased the E_MAX_ value (compound 2). Modification of the alkyl residue at R^3^ was generally better tolerated and had the potential to tune the Slack activity and pharmacological properties. Derivatives with aliphatic hydroxy groups exhibited good efficacy, with acceptable EC_50_ values (compound 4). Finally, an ionizable carboxyl group could be incorporated only at a specific linker length (compound 6), because both the addition and removal of another ethoxy unit resulted in a complete loss of activity. The combination of advantageous structural motifs did not result in a cumulative gain of activity, as compounds 7 and 8 showed improved efficacy but worse potency.

Next, we verified Slack activation by the new compounds using whole-cell patch-clamp recordings in HEK-Slack cells. In each series of experiments, compounds were added to the external solution at a concentration of 50 µM, and loxapine at a concentration of 50 µM was used as a positive control whereas vehicle (0.03% DMSO) served as a solvent control. At a holding potential of −70 mV, a series of 500 ms-long test pulses ranging from −120 to +120 mV in intervals of 20 mV were applied.^[16]^ Our patch-clamp recordings revealed that the developed Slack activators (except compound 2) significantly increased the I_K_ amplitude compared to the vehicle (Figure 1B and Figure S4, Supporting Information). At a voltage of +80 mV, the current densities of the compounds increased with a range between 2.0 to 14.8 fold compared to the baseline (Figure 1B). Notably, the efficacy of our compounds in patch-clamp recordings (i.e., the fold increase in current density) markedly correlated with their efficacy in the FluxOR assay (Figure 1C). Together, our FluxOR assay and patch-clamp analyses in HEK-Slack cells indicated that these new compounds activate Slack *in vitro*.

### 2.2. *In vitro* characterization of new Slack activators

To explore whether the compounds affected general cell functions, we performed a live-cell high-content assay that allowed for the simultaneous investigation of cellular viability, compound precipitation, and maintenance of membrane integrity and mitochondrial mass.^[25]^ HEK293 cells were stained with multiple dyes and incubated with the new compounds or loxapine at different concentrations (0.1–100 µM), and the fluorescence and cellular shape were analyzed after 6 h. We found that none of the compounds induced a relevant cytotoxic effect or affected membrane integrity, and compound precipitation was only detected for compound 1 at 100 µM whereas an increase in mitochondrial mass by more than 50% was detected only for compound 8 at 100 µM (Figure S5A–D, Supporting Information). Similarly, low cytotoxicity was observed in HEK-Slack cells, which were incubated with the compounds and analyzed by brightfield microscopy after 24 h (Figure S5E, Supporting Information). Together, these data suggest that general cell functions are not affected by any of our new compounds at concentrations up to 50 µM.

We next assessed the metabolic stability of these compounds (10 µM) in rat liver microsomes (Figure 2A). In this assay, loxapine was metabolized rapidly, with only 6% of the drug remaining unmodified after 15 min. The known metabolites of loxapine include amoxapine, 7-OH-loxapine, 8-OH-loxapine, and loxapine N-oxide.^[26]^ Therefore, we assumed that the occupation of metabolically labile sites at the aromatic core and the reduction in electron density at the oxidizable nitrogen atom of the piperazine ring might increase metabolic stability. Indeed, both the introduction of fluorine at R^2^ and the introduction of an electron-withdrawing carboxylic acid linker at R^3^ improved metabolic stability. The substitution of chlorine with a trifluoromethyl group at R^1^ was also beneficial. In contrast, homopiperazine and hydroxy linkers were metabolically labile. Compounds 6 and 7 exhibited the highest metabolic stabilities *in vitro*.

**Figure 2.**
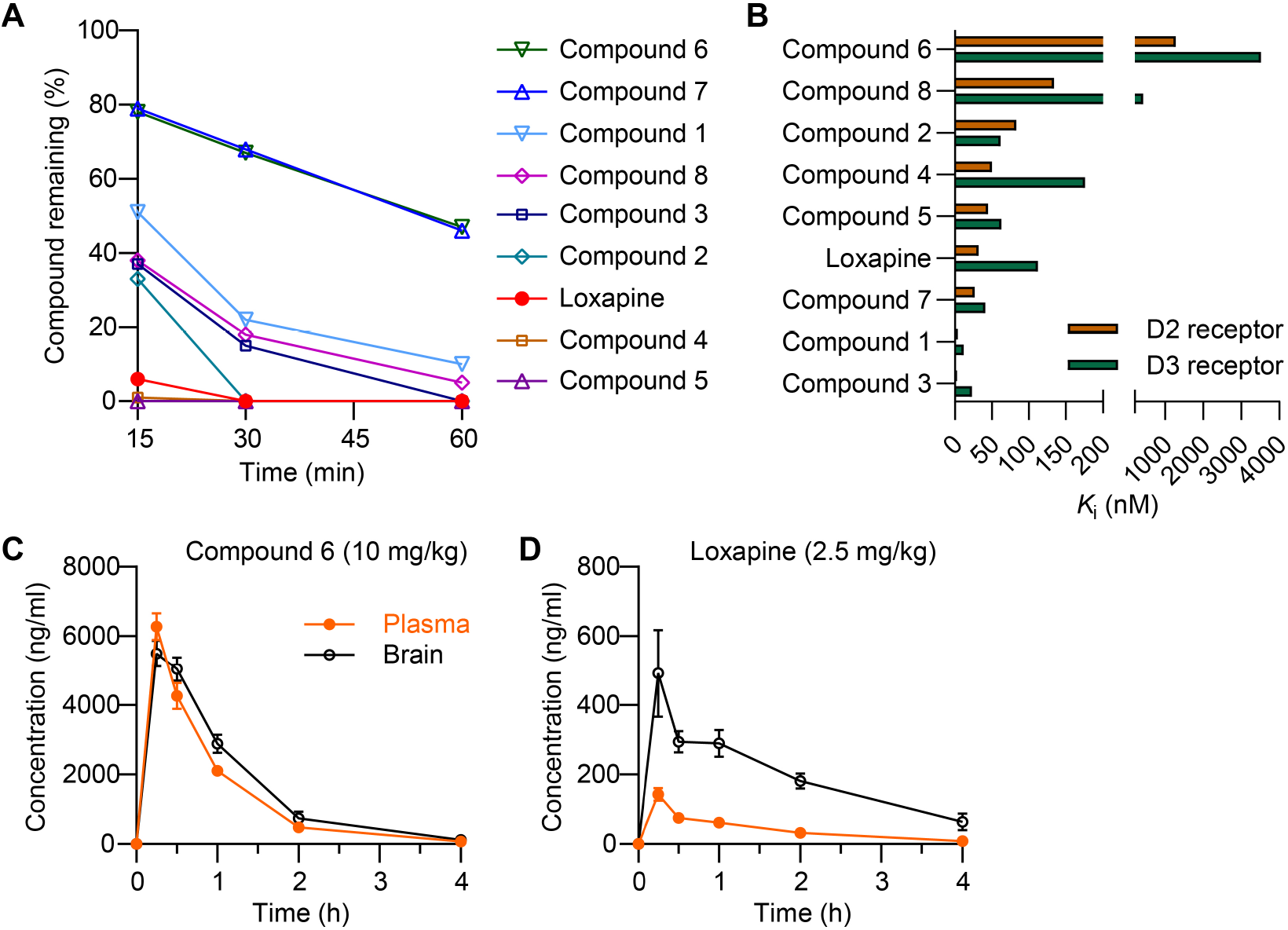
*In vitro* characterization of the new compounds and pharmacokinetics of compound 6. A) Phase I metabolism of our new compounds was assessed using a microsomal stability assay in rat liver microsomes. The percentage of compound remaining was measured 15, 30 and 60 min after compound exposure (10 µM). Data are expressed as the mean ± SEM of remaining compound from three independent experiments. B) Affinity at D_2_ and D_3_ receptors of our new compounds was determined in a [^3^H]-Spiperone competition binding assay using a cell membrane preparation of CHO cells stably expressing the human D_2S_ and D_3_ receptor. Data represent the mean; the 95% confidence interval and *n* number are presented in Table S1 (Supporting Information). Data in (A) and (B) show that compound 6 has a high microsomal stability and a low affinity to D_2_ and D_3_ receptors. C,D) Pharmacokinetic properties of compound 6 in comparison to loxapine. Animals were i.p. injected with 10 mg/kg compound 6 (C) or 2.5 mg/kg loxapine (D) and plasma and brain levels were measured at the indicated time points by LC-MS analysis. Data are presented as mean ± SEM of 3 mice per group. Additional pharmacokinetic parameters are presented in Table S2 (Supporting Information).

Because some of the most limiting adverse effects of loxapine are mediated by the antagonism of dopamine receptors, we aimed to develop derivatives with decreased dopamine receptor affinity. Therefore, we determined the binding affinities of our compounds to D_2_ and D_3_ dopamine receptors using a [^3^H]-spiperone displacement assay (Figure 2B and Table S1, Supporting Information). As expected, loxapine displayed pronounced binding to the D_2_ and D_3_ receptors, with *K*_i_ values of 32 and 112 nM, respectively. Generally, the affinity was further promoted by the trifluoromethyl group at R^1^. Derivatives with a homopiperazine ring or hydroxy linker had a slightly lower affinity. Importantly, compound 6 exhibited a ∼40-fold lower affinity against D_2_ and a ∼31-fold lower affinity against D_3_ as compared to loxapine. These findings prompted us to focus on compound 6 for the remainder of this study.

### 2.3. Target binding and off-target profile of compound 6

To confirm that the cellular effect of compound 6 was mediated by its direct interaction with Slack, ligand binding on purified human Slack (Figure S6A, Supporting Information) was assessed by differential scanning fluorimetry (DSF), thereby evaluating a ligand-dependent shift of the protein melting temperature (*T_m_*). The cysteine-reacting dye 7-diethylamino-3-(4-maleimidophenyl)-4-methylcoumarin (CPM), which binds to cysteine residues that are accessible during protein unfolding, was used as an indicator for protein unfolding. In the absence of ligand, Slack had a *T_m_* of 30 °C. In presence of a Slack inhibitor (compound 31 from ref^[27]^), known to directly interact with the protein and therefore used as a positive control, *T_m_* saturated at 47 °C at a ligand concentration of 100 µM (Figure S6B, Supporting Information), indicating a notable stabilization of Slack. Similarly, protein unfolding with compound 6 and loxapine reached its maximum at 100 µM ligand concentration with *T_m_*s of 39 and 37 °C, respectively, reflecting an increase in *T_m_* of 9 and 7 °C, respectively. These results confirmed that compound 6 and loxapine acted as direct ligands on Slack (Figure S6B, Supporting Information).

As loxapine is a first-generation antipsychotic with a substantial binding affinity for many receptors,^[26]^ we next explored the pharmacological profile of compound 6 and loxapine *in vitro* using a SafetyScreen44 panel (Eurofins), which measures the interaction of compounds with 44 targets. In this panel, loxapine (10 µM) showed substantial binding inhibition (higher than 50%) for 17 out of 44 targets including adrenergic (α_1A_ and α_2A_), dopamine (D_1_ and D_2S_), histamine (H_1’_and H_2_), muscarinic (M_1_, M_2_ and M_3_), and serotonin (5-HT_1A_, 5-HT_1B_, 5-HT_2A_, 5-HT_2B_ and 5-HT_3_) receptors; Na^+^ channel; norepinephrine transporter (NET); and serotonin transporter (SET) (Figure S7, Supporting Information), thereby confirming the “dirty” nature of first-generation antipsychotics. Of note, considerably less off-target activities were observed for compound 6 (10 µM), which exhibited substantial binding to 8 targets (α_1A_, D_1_, D_2S_, H_1_, 5-HT_2A_, 5-HT_2B_, NET, and SET). These data suggested an improved pharmacological profile of compound 6 compared to that of loxapine. Furthermore, compound 6 (like loxapine) did not measurably bind to human ether-à-go-go (hERG) or voltage-gated (K_V_) potassium channels in the off-target screen (Figure S7, Supporting Information), suggesting that this compound does not act as an unspecific modulator of potassium channels.

### 2.4. Compound 6 profoundly inhibits itch-related behavior in mice

To prepare for *in vivo* studies, we investigated the pharmacokinetic properties of compound 6. Following intraperitoneal (i.p.) administration of compound 6 (10 mg/kg) in mice, plasma and brain levels were determined over 4 h using LC-MS. We found that compound 6 concentrations decayed with first-order kinetics in the plasma and brain, with a half-life of approximately 35 min, and readily crossed the blood-brain barrier (Figure 2C and Table S2, Supporting Information). No obvious impairment of mouse behavior was observed after the administration of compound 6 at this dose. For comparison we determined the plasma and brain loxapine levels. As i.p. administration of 10 or 5 mg/kg loxapine caused obvious signs of toxicity (i.e., considerably reduced activity), loxapine was administered at a dose of 2.5 mg/kg. Relatively low concentrations of loxapine were detected in the plasma (Figure 2D and Table S2, Supporting Information), which was consistent with its rapid metabolism (Figure 2A).^[28]^ Moreover, loxapine had an especially high brain exposure (ratio_brain/plasma_ = 5.12) that is in accordance with its activity on dopamine receptors in the central nervous system (CNS), unlike compound 6 (Figure 2B). Collectively, the pharmacological and pharmacokinetic profiles of compound 6 make it an excellent molecule for *in vivo* pharmacodynamic studies.

The *in vivo* efficacy of compound 6 was analyzed in mice of both sexes, and no significant sex-related differences were observed in the behavioral experiments (Figures S8 and S9, Supporting Information). As a prerequisite for interpreting itch behavior experiments, we first tested motor function after systemic dosing. Treatment with compound 6 (3–30 mg i.p.) did not impair motor function in the accelerating rotarod test, a standard model of motor performance (Figure 3A), or in the vertical pole test, which assesses basal ganglia-related movement disorders (Figure 3B). By contrast, loxapine at a dose of ≥ 0.39 mg/kg markedly reduced the ability of the mice to perform in both models (Figure 3A,B), which is in line with its high CNS penetration and dopamine receptor affinity, thereby confirming earlier reports.^[24]^

**Figure 3.**
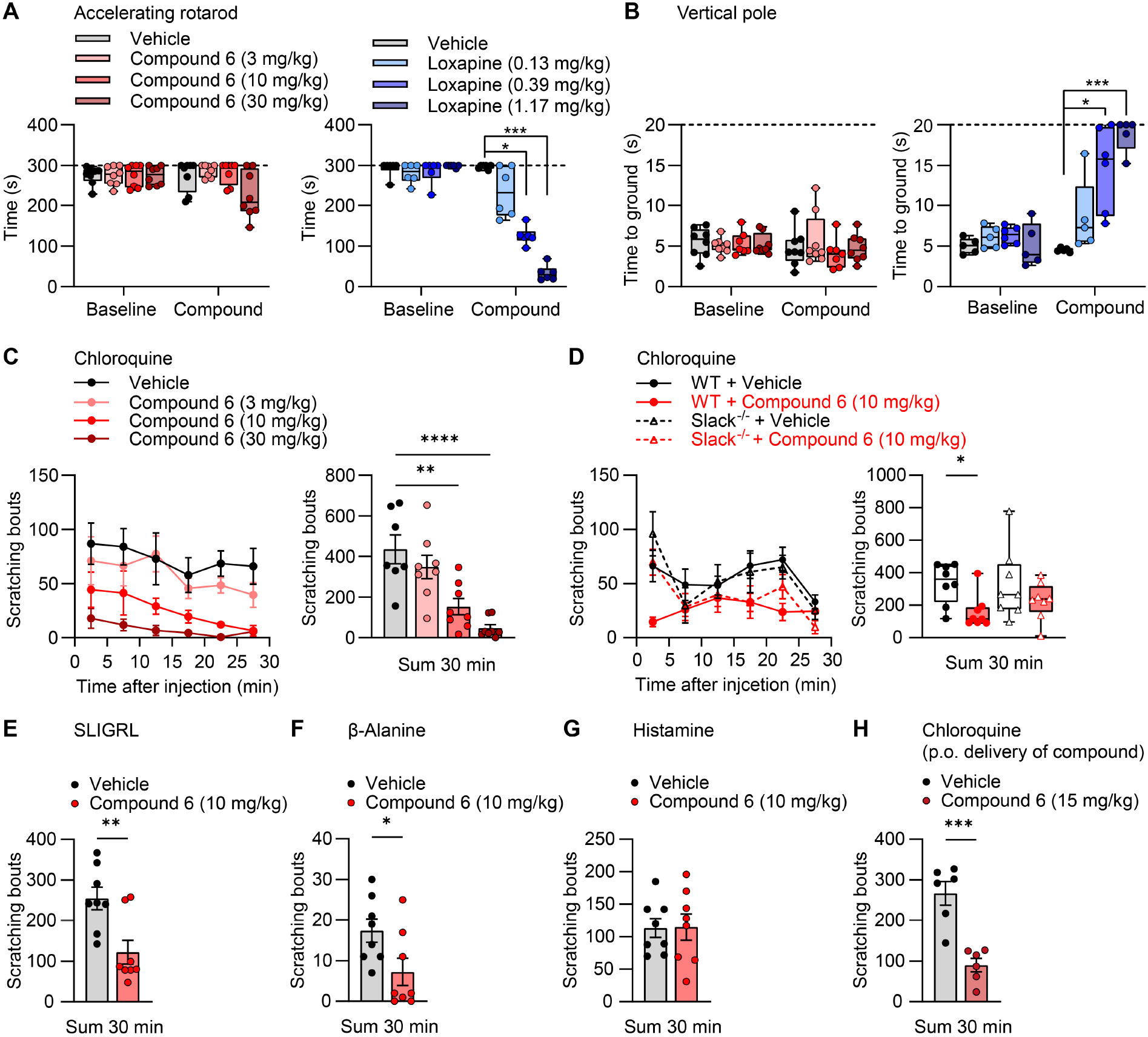
Compound 6 inhibits histamine-independent acute itch behavior. A,B) Motor function. Compound 6, loxapine or vehicle (0.9% NaCl with 10% 2-hydroxypropyl-β-cyclodextrine) were i.p. administered and an accelerating rotarod test (A) followed by a vertical pole test (B) were performed 15 min thereafter. Data show that compound 6 did not affect the time spent on the rotarod or the vertical pole, whereas loxapine dose-dependently inhibited the motor coordination in both models. Box-and-whisker plots represent maximum and minimum values, and the box shows the first, second (median) and third quartile values. Dotted lines indicate cutoff times (*n* = 6–8). \**P* < 0.05, \*\*\**P* < 0.001, Kruskal-Wallis test. C,D) Chloroquine-induced itch behavior. Compound 6 or vehicle were i.p. administered. After 15 min, chloroquine was s.c. administered into the nape of the neck and the number of scratching bouts was counted over 30 min. In each panel, the time course of scratching behavior is shown on the left and the sum of scratching bouts in 30 min is presented on the right. Note that compound 6 inhibited the scratching behavior induced by chloroquine in a dose-dependent manner (C) and that compound 6 significantly inhibited the chloroquine-induced scratching behavior in WT mice but not in Slack^-/-^ littermates (D). In C, data represent the mean ± SEM (*n* = 7–8). Vehicle (veh) vs. 3 mg/kg, *P* = 0.4713; veh vs. 10 mg/kg, *P* = 0.0013; veh vs. 30 mg/kg, *P* = < 0.0001; one-way-ANOVA with Dunnett’s correction. In D, data in the time course diagram represent the mean ± SEM; data in the sum diagram are presented as box-and-whisker plots with maximum and minimum values, in which the box shows the first, second (median), and third quartile values (*n* = 8). WT (veh vs. compound 6), *P* = 0.0242; Slack^-/-^ (veh vs. compound 6), *P* > 0.9999; Kruskal-Wallis test with Dunn’s correction. E–G) Itch behavior induced by SLIGRL, β-alanine or histamine. Compound 6 significantly ameliorated the scratching behavior induced by SLIGRL (E) or β-alanine (F), but not by histamine (G). Data represent the mean ± SEM (*n* = 8). *P* = 0.0053 (E), *P* = 0.0365 (F), *P* = 0.9563 (G); unpaired Student’s *t*-test. H) Acute itch behavior in the chloroquine model was also inhibited after p.o. administration of compound 6 under similar experimental settings as in C but with p.o. delivery of compound 6 or vehicle. Data represent the mean ± SEM (*n* = 6). *P* = 0.0004; unpaired Student’s *t*-test. \**P* < 0.05, \*\**P* < 0.01, \*\*\**P* < 0.001, \*\*\*\**P* < 0.0001.

We then assessed the antipruritic efficacy of compound 6 in a model of histamine-independent itching induced by the antimalarial drug chloroquine. Importantly, we found that systemic treatment of mice with compound 6 (3, 10, or 30 mg/kg i.p.) 15 min prior to subcutaneous (s.c.) injection of chloroquine into the nape of the neck ameliorated scratching behavior in a dose-dependent manner (Figure 3C). Further experiments supported the conclusion that the antipruritic effects were mediated by Slack activation, because compound 6 did not significantly alter chloroquine-induced scratching in animals with a genetic deletion of Slack (Slack^-/-^ mice; Figure 3D). Taken together, these data suggest that treatment with compound 6 ameliorates chloroquine-induced itching by activating Slack.

Chloroquine-induced scratching in mice is driven by MrgprA3 activation in the NP2 population of sensory neurons.^[10, 29]^ To further explore the antipruritic efficacy of compound 6 in histamine-independent itch, we assessed the behavioral response after s.c. injection of the peptide Ser-Leu-Ile-Gly-Arg-Leu (SLIGRL) that evokes scratching by activation of MrgprC11,^[30]^ which is expressed in the NP2 and NP3 population.^[10]^ Similar to the chloroquine model, compound 6 (10 mg/kg i.p.) significantly inhibited SLIGRL-induced scratching behavior (Figure 3E). Moreover, compound 6 ameliorated histamine-independent itch evoked by s.c. injection of β-alanine that activates MrgprD^[31]^ in the NP1 population of sensory neurons^[10]^ (Figure 3F). In contrast, scratching induced by subcutaneous histamine was not affected by compound 6 (Figure 3G). The lack of an effect on histamine-evoked scratching was supported by the low affinity of compound 6 for histamine H_1_ receptors, which was determined using a [^3^H]-pyrilamine competition binding assay *in vitro* (Table S3, Supporting Information). In addition, the unaltered scratching behavior in the histamine model provided further evidence that the effects of compound 6 observed in the histamine-independent models did not result from impaired motor function. We also investigated the efficacy of compound 6 after peroral (p.o.) delivery. Consistent with the relatively high exposure after oral dosing (Table S2, Supporting Information), this molecule also conferred a profound antipruritic effect when delivered at a dose of 15 mg/kg p.o. in the chloroquine itch model (Figure 3H).

We next assessed the efficacy of compound 6 in the treatment of chronic itching. In a model of allergic contact dermatitis,^[32]^ painting the nape of the neck with the hapten 2,4-dinitrofluorobenzene (DNFB) twice two weeks apart (Figure 4A) resulted in persistent itching behavior reflected by scratching the painted area and head-shaking, which was significantly correlated with scratching (Figure S9E, Supporting Information). Interestingly, compound 6 administered i.p. after the second DNFB exposure significantly reduced the number of scratching bouts and head shakes compared to vehicle-treated animals (Figure 4B,C). Similarly, in the MC903 model of skin inflammation (Figure 4D), which has some characteristics of allergic contact dermatitis and atopic dermatitis,^[33, 34]^ compound 6 significantly ameliorated scratching and head shaking (Figure 4E,F). These findings are important because they indicate that Slack activators can reduce existing itching.

**Figure 4.**
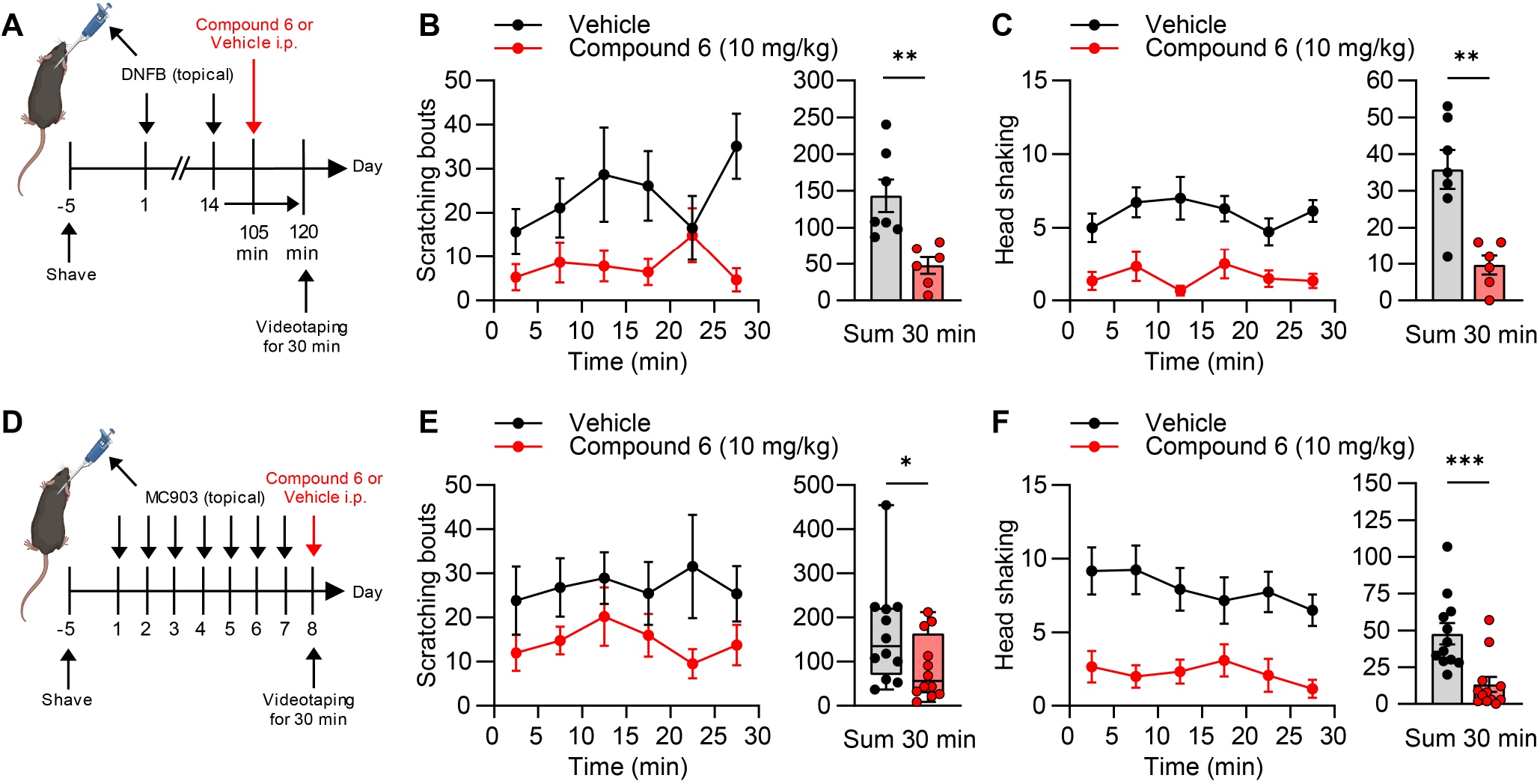
Compound 6 inhibits persistent itch behavior. A–C) Efficacy of compound 6 in the DNFB model of persistent itch. A) Experimental diagram showing the induction of spontaneous itch behavior by topical application of 2,4-dinitrofluorobenzene (DNFB) to the nape of the neck (twice 14 days apart), i.p. drug delivery 105 min after the second DNFB application, and videotaping 15 min thereafter. Compound 6 significantly reduced the number of scratching bouts (B) and the number of head shakes (C) as compared to vehicle. Data are presented as mean ± SEM (*n* = 6–7). *P* = 0.004, unpaired Student’s *t*-test (B); *P* = 0.0014, unpaired Student’s *t*-test (C). D–F) Efficacy of compound 6 in the MC903 model of persistent itch. D) Experimental diagram showing the induction of spontaneous scratching by topical application of MC903 to the nape of the neck (once daily over 7 days), i.p. drug delivery at day 8, and videotaping 15 min thereafter. In comparison to the vehicle, compound 6 significantly reduced the number of scratching bouts (E) and the number of head shakes (F). In E, box-and-whisker plots represent maximum and minimum values, and the box shows the first, second (median) and third quartile values. All other data are presented as mean ± SEM (*n* = 12). *P* = 0.0319, Mann-Whitney test (E); *P* = 0.0008, unpaired Student’s *t*-test (F);. \**P* < 0.05, \*\**P* < 0.01, \*\*\**P* < 0.001.

Finally, compound 6 was screened for potential side effects *in vivo*. In pulse oximetry measurements of conscious, freely moving mice, i.p. administration of compound 6 (10 mg/kg) did not affect the heart rate (Figure S10A, Supporting Information) or breathing rate (Figure S10B, Supporting Information) during an observation period of 30 min. In contrast, both parameters were reduced after i.p. administration of morphine (10 mg/kg), which was used as a positive control in this experiment. Because Slack-positive sensory neurons are polymodal and involved in the sensation of mechanical and heat stimuli,^[35]^ we also tested the response to mechanical (von Frey filament test) and thermal (Hargreaves test) stimulation of the hindpaws after i.p. administration of compound 6 (10 mg/kg). Neither mechanical nor thermal sensitivity was altered by compound 6 compared to the vehicle control (Figure S10C,D, Supporting Information). Taken together, these results highlight that compound 6 effectively inhibited the itch behavior of mice in multiple models at a well-tolerated dose.

### 2.5. Compound 6 inhibits itch-sensitive sensory neurons

To determine whether the antipruritic effect of compound 6 occurs directly at the neuronal level, we used whole-cell patch-clamp electrophysiology to measure the excitability of sensory neurons. In order to simulate a pathological state, we incubated dorsal root ganglia (DRG) neuronal cultures with an “inflammatory soup” (histamine: 10 µM, PGE_2_: 10 μM, serotonin: 10 μM, bradykinin: 10 μM)^[36]^ overnight and performed recordings on cells that bind isolectin B4, a marker of non-peptidergic C-fiber neurons including the itch-sensitive neuronal populations (NP1–NP3)^[10]^ in mice. Neurons from these cultures displayed pronounced hyperexcitability, as indicated by spontaneous action potential (AP) firing after current injection (200–950 pA; Figure 5A). Notably, application of compound 6 (50 µM), but not of vehicle, caused a profound decrease in the number of AP fired (Figure 5A,B). Neither compound 6 nor the vehicle significantly affected the resting membrane potential (RMP; Figure 5B). Furthermore, after injection of small currents (0–220 pA), the rheobase (i.e., the amount of current required to generate an AP) and the AP duration at 90% repolarization (APD90) were markedly increased in presence of compound 6, whereas the AP amplitude was considerably lower (Figure 5C,D). No significant alterations in the rheobase, AP duration, or AP amplitude were observed in the in presence of vehicle (Figure 5C,D). These findings confirmed that compound 6 inhibited itch-sensitive sensory neurons in mice.

**Figure 5.**
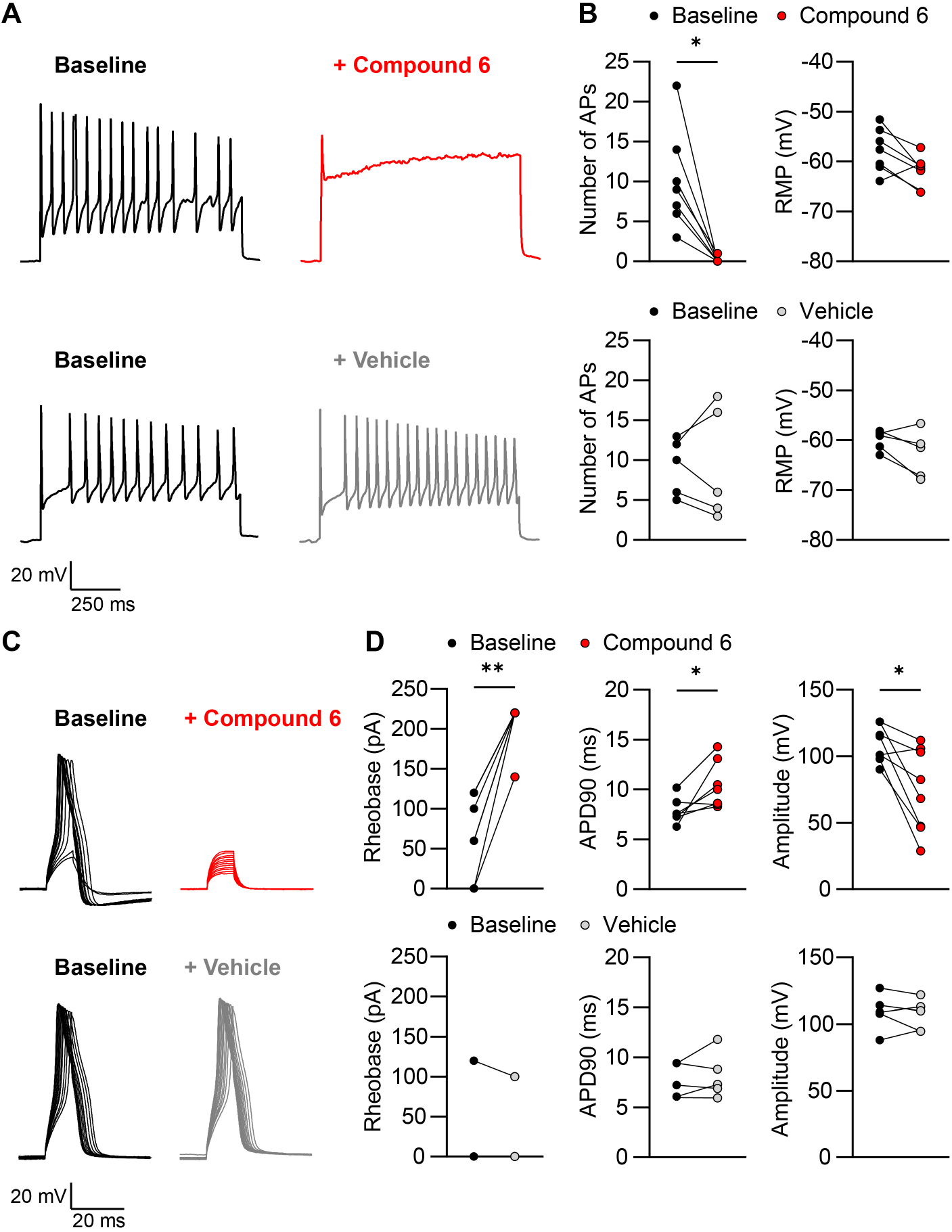
Compound 6 reduces neuronal excitability of itch-sensitive sensory neurons. DRG neurons were incubated overnight with an “inflammatory soup” followed by whole cell current-clamp recordings on IB4-binding neurons. A) Representative recording from a neuron showing action potential (AP) firing in response to current injections (200–950 pA at 150 pA intervals, 1000-ms duration) at baseline and in the presence of compound 6 (50 µM) or vehicle. B) Group data show that compound 6 significantly blocked the number of APs compared to baseline but did not significantly affect the resting membrane potential (RMP). Baseline vs. compound 6: number of APs, *P* = 0.0156, Wilcoxon test (*n* = 7); RMP, *P* = 0.2812, paired Student’s *t*-test (*n* = 7). Baseline vs. vehicle: number of APs, *P* = 0.9193, paired Student’s *t*-test (*n* = 5); RMP, *P* = 0.1226, paired Student’s *t*-test (*n* = 5). C) Recordings from DRG neurons showing single APs evoked by injections of small currents (0–220 pA at 20 pA intervals, 10-ms duration) at baseline and in the presence of compound 6 (50 µM) or vehicle. D) Group data indicate that compound 6, but not vehicle, significantly increased the rheobase (amount of current required to generate an AP) and the AP duration at 90% repolarization (APD90), as well as reduced the AP amplitude. Baseline vs. compound 6: rheobase, *P* = 0.0078, Wilcoxon test (*n* = 8); APD90, *P* = 0.0257, paired Student’s *t*-test (*n* = 7); amplitude, *P* = 0.0103, paired Student’s *t*-test (*n* = 8). Baseline vs. vehicle: rheobase, *P* > 0.9999, Wilcoxon test (*n* = 5); APD90, *P* = 0.4123, paired Student’s *t*-test (*n* = 5); amplitude, *P* = 0.5907, paired Student’s *t*-test (*n* = 5). Data in B and D represent the individual responses to compound 6 or vehicle compared to baseline. \**P* < 0.05, \*\**P* < 0.01.

## 3. Discussion

The research presented here identifies several new compounds that can activate Slack channels. Our finding that the lead molecule, compound 6, effectively inhibited scratching elicited by multiple histamine-independent pruritogens in mice established Slack activation as a novel pharmacological strategy for the treatment of antihistamine-resistant itch.

Our aim to develop Slack-activating compounds for the treatment of itch was based on the finding that Slack expression is enriched in all subtypes of itch-sensing sensory neurons (Figure S1, Supporting Information) and that Slack controls the bursting and adaptation of action potential firing rates, thereby reducing the neuronal activity of sensory neurons.^[24, 37–39]^ We started compound optimization from an approved drug, which is considered to be a highly favorable approach known as selective optimization of side activities (SOSA).^[40, 41]^ Loxapine has been previously discovered as a Slack activator,^[21]^ however the pharmacodynamic properties of loxapine clearly limit its applicability in patients.^[23]^ Moreover, the lack of selectivity toward key aminergic receptors prevents the applicability of loxapine as a reliable pharmacological tool to validate Slack as a therapeutic target. In the case of the loxapine-derivative compound 6, the replacement of the methyl residue by the negatively charged ethoxyacetic acid moiety led to a substantial reduction in off-target interactions, while the efficacy and potency of Slack activation were similar to those of loxapine. Overall, these properties qualify compound 6 as a valuable pharmacological tool for investigating the effects of Slack activation *in vivo*. Indeed, the profound inhibition of scratching behavior in mice after administration of compound 6 (10–30 mg/kg) suggests that this compound sufficiently activates Slack *in vivo*, although its FluxOR EC_50_ value (30 µM) is relatively high. When interpreting the FluxOR EC_50_ values, it should be considered that these experiments were performed using an assay buffer containing 2 mM Ca^2+^. A previous study revealed that Ca^2+^ inhibits Slack activity with an IC_50_ of 0.26 mM when applied to the intracellular side of the membrane in inside-out patches.^[15]^ Although highly speculative, the EC_50_ values measured in our FluxOR assay may have been affected by Ca^2+^-mediated Slack inhibition in this *in vitro* approach. Nevertheless, the assumption that Slack is the primary target mediating antipruritic activity *in vivo* is supported by the observation that the action of compound 6 was largely abolished in Slack^-/-^ mice. Based on the hits in the off-target screen, it might be possible that the binding of compound 6 to 5HT_2A_ receptors also affects its antipruritic efficacy because serotonin causes itching via the activation of several 5-HT receptors, most commonly 5-HT_2_ (ref^[42]^). 5-HT_2A_ (but not 5-HT_2B_) is expressed in sensory neurons.^[43]^ It must be taken into account, however, that in our off-target screen, the compounds were incubated at only one concentration (10 µM). Determination of receptor affinities will be important in future studies to further explore the extent of off-target interactions.

We found that compound 6 inhibited the scratching behavior of mice in the chloroquine, SLIGRL and β-alanine model of acute non-histaminergic itch and in the DNFB and MC903 model of chronic itch. Furthermore, compound 6 profoundly inhibited the excitability of IB4-binding sensory neurons in mice. The strong efficacy of compound 6 in mice, together with the prominent expression of Slack in pruritogen receptor-enriched sensory neurons in humans (Figure S1D, Supporting Information), suggests that Slack activators may also be effective for the treatment of itch in humans. However, it should be considered that the mechanisms underlying itch sensation and processing in humans are poorly understood. Moreover, there are substantial differences in the expression of itch-relevant genes between species. For example, mice express 27 Mrgprs, whereas humans express eight Mrgprs (MRGPRX1–X4 and MRGPRD–G). MRGPRX1, -X4, and -D are expressed in the human DRG and trigeminal ganglia and have been implicated in non-histaminergic itch as receptors for pruritic compounds.^[44]^ Further studies are needed to elucidate the functional role of Slack in the human itch sensation.

Although we did not extensively evaluate the side effects, we found that motor coordination (Figure 3A,B), heart rate or breathing rate (Figure S10A,B, Supporting Information), and the sensing of mechanical or thermal stimuli (Figure S10C,D, Supporting Information) were not affected by the administration of compound 6 at an effective antipruritic dose. With regard to the safety profile, it should be considered that Slack is expressed in various neuronal populations in the CNS, and gain-of-function mutations of *KCNT1* have been associated with epileptic encephalopathies in humans.^[45]^ Limited information exists concerning the incidence of seizures during loxapine treatment. The overall risk for loxapine-induced seizures seems to be low at therapeutic doses,^[22, 26]^ but it should be used with caution in patients with a history of convulsive disorders. In our behavioral studies in mice, we did not observe any overt signs of seizures after the systemic administration of compound 6 at doses up to 30 mg/kg. However, the epileptogenic potential of brain-penetrant Slack activators such as compound 6 should be carefully assessed in the future.

In summary, compound 6 represents a new family of Slack activators. Its profound efficacy in various itch models illustrates its potential as an antipruritic drug for the treatment of histamine-independent and chronic itch.

## 4. Experimental Section

### 4.1. Chemicals/Drugs

Loxapine, pregabalin, chloroquine, SLIGRL-NH_2_, β-alanine, histamine, PGE_2_, serotonin, and bradykinin were purchased from Sigma-Aldrich. All indicated concentrations refer to those of pure substances. Reagents and solvents for the synthesis of loxapine derivatives were obtained from Acros Organics (Gel, Belgium), Alfa Aesar GmbH & Co KG (Karlsruhe, Germany), BLDPharm Inc. (NuiNan, China), Fluorochem Ltd. (Hadfield, UK), Sigma-Aldrich (Munich, Germany), and TCI Europe N.V. (Zwijndrecht, Belgium).

### 4.2. Design, preparation and analytical characterization of loxapine derivatives

Reactions were carried out in an argon atmosphere. NMR spectra were recorded on Bruker DPX250, Bruker Avance 300, Bruker Avance 400, or Bruker Avance 500 (Bruker, Karlsruhe, Germany) operating at ambient temperature. Proton spectra were recorded in CDCl_3_ or DMSO-d_6_, and ^1^H NMR chemical shifts were referenced to the residual signals of CHCl_3_ (at δ = 7.26 ppm) and DMSO-d_5_ (at δ = 2.50 ppm). ^13^C NMR chemical shifts were referenced against the central line of the solvent signal (for CHCl_3_ at δ = 77.16 ppm and for DMSO-d_6_ at δ = 39.52 ppm). Chemical shifts are given on δ scale (ppm). The coupling constants (J) are given in hertz (Hz). The multiplicities are as follows: s (singlet), d (doublet), t (triplet), q (quartet), quint or m (multiplet). ESI-MS was performed with an LCMS-2020 from Shimadzu and HRMS with a MALDI Orbitrap XL (Thermo Scientific). TLC was carried out on silica gel plates from Marcherey-Nagel (ALUGRAM^®^) and visualized with a UV lamp (254 nm and/or 366 nm). Purification of products was performed by flash chromatography using puriFlash XS420 and Silica HP 30 µm columns as stationary phase (Interchim, Montluçon, France). Analytical and semi-preparative HPLC was conducted using Shimadzu Prominence with an SPD20A UV/Vis detector (Shimadzu, Duisburg, Germany). Stationary phases were Luna 10 µm 100 Å, C18(2) (250 x 4.6 mm), and Luna 10 µm 100 Å, C18(2) (250 x 21.20 mm), from Phenomenex (Aschaffenburg, Germany), and the eluent was a mixture of ACN and aqueous formic acid solution (0.1%). Flow rates were set to 1 mL/min and 21 mL/min. All compounds tested displayed a purity of > 95% (254 nm).

### 4.3. Cell cultures

HEK293 cells stably transfected with human *KCNT1* (herein referred to as HEK-Slack cells; SB-HEK-KCa4.1; SB Drug Discovery, Lanarkshire, UK) were maintained in Dulbecco’s modified Eagle’s medium-Glutamax with 10% fetal calf serum and 1% penicillin/streptomycin, supplemented with 0.6 mg/mL G-418 (all from Gibco/Thermo Fisher Scientific) in 5% CO_2_ at 37 °C. Cells were passaged every 4–5 days from P11 to P35 depending on their confluence.

### 4.4. FluxOR assay

To determine the Slack-activating efficacy and potency of the novel compounds, a commercial FluxOR^TM^ potassium ion channel assay (#10017; Invitrogen) was used. HEK-Slack cells were plated at a density of 50,000 cells per well on a poly-D-lysine coated (75 μg/mL, Sigma Aldrich) 96-well, black-walled microplate with clear bottom (#655090; Greiner Bio-One) in DMEM (100 µL/well) containing 10% FBS and 1% penicillin/streptomycin 24 h prior to assaying. In the initial experiments shown in Figure S3 (Supporting Information), HEK-Slack cells were plated at a density of 100,000 cells/well 48 h prior to the assay. On the day of the experiment, the cells were loaded with FluxOR^TM^ dye according to the manufacturer’s protocol. A Na^+^-free assay buffer, adjusted to pH 7.4 with KOH, containing 140 mM choline chloride, 5 mM KCl, 2 mM CaCl_2_, 2 mM MgSO_4_, 10 mM HEPES, and 5.55 mM glucose was used. To reduce background fluorescence, all assay components were complemented with 10% BackDrop^TM^ Background Suppressor (#B10511; Invitrogen).

All compounds were prepared as a 333 mM stock in DMSO and diluted at the experimental day to a final concentration of 100, 50, 25, 12.5, 6.25, 3.125, or 1.5625 μM in an assay buffer with DMSO adjusted to 0.03%. Before treatment, the cells were washed, and the medium was exchanged for the assay buffer. In a total volume of 90 µL/well, the cells were then incubated for 30 min at 37 °C with compounds at the indicated concentrations, or 0.03% DMSO alone as vehicle. A positive control with 50 μM loxapine was always conducted in parallel. Fluorescence was measured using an Infinite M200 microplate reader (Tecan) with excitation at 485 nm (9 nm bandwidth) and emission at 525 nm (20 nm bandwidth) in the bottom read mode. First, the baseline fluorescence of the individual wells was established. Thereafter cells were stimulated with 20 µL Tl_2_SO_4_ (5mM) solution injected at 100 µL/sec, and fluorescence was further recorded for 100 s. Assays were run in triplicate. For all time points, the ratio of fluorescence to baseline fluorescence was calculated first for individual wells, and the average was then used.

Dose-response curves comprising six concentrations were obtained in triplets using fluorescence/fluorescence_Baseline_ ratios at the 100^th^ s. Values were calculated relative to loxapine in each experiment as follows:100 × ([F/F_B_(Compound) - F/F_B_(Vehicle_mean_)] / [(F/F_B_(Loxapine) - F/F_B_(Vehicle_mean_]). To generate EC_50_ and E_MAX_ values, a standard, logistic, nonlinear regression analysis with GraphPad Prism 9.0 was used.

### 4.5. Patch-clamp recordings

#### 4.5.1. HEK-Slack cells

HEK-Slack cells were plated onto poly-D-lysine-coated (100 μg/mL, Sigma Aldrich) coverslips 1 d before experiments and cultured in DMEM containing 10% FBS and 1% penicillin/streptomycin in 5% CO_2_ at 37 °C. Whole-cell voltage-clamp recordings were acquired using an EPC 9 amplifier combined with Patchmaster software (HEKA Electronics, Lambrecht/Pfalz, Germany). Currents were sampled at 20 kHz and filtered at 5 kHz. Data analysis was performed using the Fitmaster software (HEKA Electronics). Membrane potential was held at −70 mV and outward K^+^ current (I_K_) was evoked by depolarizing steps (500 ms duration) ranging from −120 to +120 mV in increments of 20 mV. The pipette solution contained 140 mM KCl, 2 mM MgCl_2_, 5 mM EGTA, and 10 mM HEPES and was adjusted to pH 7.4 with KOH. The extracellular solution contained 140 mM NaCl, 5 mM KCl, 2 mM CaCl_2_, 2 mM MgCl_2_, and 10 mM HEPES and was adjusted to pH 7.4 with NaOH. The osmolarity of all the solutions was adjusted to 290–300 mOsmol/L using glucose. Patch pipettes had a resistance of 6–8 MΩ and were obtained from borosilicate glass capillaries (Science Products) using a conventional puller (DMZ-Universal Puller, Zeitz Instruments).

After baseline measurements, new compounds or loxapine solved in external solution containing 0.03% DMSO (each with a final concentration of 50 µM) were added to the bath without a continuous perfusion, and K^+^ currents were measured within 5 min. The fold-increase values in the patch-clamp experiments were determined by calculating the baseline current density relative to the current density after compound application.

#### 4.5.2. DRG neurons

A primary cell culture of DRG neurons was prepared and stimulated with an inflammatory soup overnight to induce hyperexcitability. Briefly, naïve C57BL/6N mice (aged 4–8 weeks) were sacrificed by CO_2_ inhalation, and lumbar (L1–L5) DRGs were transferred to a hanks’ balanced salt solution (Gibco, Thermo Fisher Scientific). After incubation with 500 U/mL collagenase IV and 2.5 U/mL dispase II (both from Sigma Aldrich) for 60 min, including carefully shaking every 20 min, DRGs were washed and gently triturated twice with a fire-polished Pasteur pipette in neurobasal medium (Gibco, Thermo Fisher Scientific) including 10% FBS and 0.5 mM GlutMax (Gibco, Thermo Fisher Scientific). Dissociated DRGs were seeded onto poly-D-lysine-coated (100 μg/mL) coverslips and cultured in neurobasal medium supplemented with 2% B27 (Gibco, Thermo Fisher Scientific), 1% penicillin/streptomycin, and 0.5 mM GlutMax in 5% CO_2_ at 37 °C. In order to increase neuronal excitability, DRG cultures were incubated with an inflammatory soup (histamine:10 μM, PGE_2_:10 μM, serotonin:10 μM, and bradykinin:10 μM)^[36]^ overnight. Before recording, DRG neurons were preincubated with 10 μg/mL FITC-conjugated IB4 (Sigma-Aldrich) for 5–10 min to select Slack-expressing neurons. Whole-cell current clamp recordings were obtained using an EPC 9 amplifier combined with Patchmaster software (HEKA Electronics, Lambrecht/Pfalz, Germany). General settings, pipettes, and extracellular solutions were used as described above. Evoked action potentials (APs) were elicited by 10 ms current injections starting at 0 pA in 20 pA increments to determine electrophysiological parameters. AP firing was induced by depolarizing current pulses (200–950 pA at 150 pA intervals, 1,000 ms duration). Recordings were performed at baseline and following a 1 min incubation with 50 µM compound 6 or vehicle (external solution containing 0.03% DMSO). The number of APs before and after the compound application was counted at the same current-injection step.

### 4.6. Cytotoxicity multiplex assay

#### 4.6.1. HEK293T cells

To assess cell viability, a live cell high-content screen was performed in HEK293T cells as described previously.^[25]^ In brief, HEK293T cells (ATCC® CRL-1573™) were cultured in DMEM plus L-glutamine (high glucose) supplemented by 10% FBS (Gibco) and penicillin/streptomycin (Gibco). Cells were seeded at a density of 1,500 cells/well in 384 well plates in culture medium (cell culture microplate, PS, f-bottom, µClear®, 781091; Greiner). Cells were stained simultaneously with 60 nM Hoechst33342 (Thermo Scientific), 75 nM Mitotracker red (Invitrogen), 0.3 µL/well annexin V Alexa Fluor 680 conjugate (Invitrogen), and 25 nL /well BioTracker™ 488 green microtubule cytoskeleton dye (EMD Millipore). The fluorescence and cellular shape were measured after 6 h of compound exposure using a CQ1 high-content confocal microscope (Yokogawa). The following setup parameters were used for image acquisition: Ex 405 nm/Em 447/60 nm, 500 ms, 50%; Ex 561 nm/Em 617/73 nm, 100 ms, 40%; Ex 488/Em 525/50 nm, 50 ms, 40%; bright field, 300 ms, 100% transmission, one centered field per well, 7 z stacks per well with 55 µm spacing. Images were analyzed using the CellPathfinder software (Yokogawa). Cells were detected and gated in different cell populations as described previously^[46]^ using a machine learning based algorithm. Briefly, compound precipitation was detected as Hoechst high intensity objects. Cell viability was calculated based on the normal cell population (cells that did not show Hoechst high intensity objects). Healthy cells were further gated into phenotypic subpopulations (membrane integrity and mitochondrial mass). The compounds were tested in duplicates. Cell counts of different populations were normalized against cells treated with 0.1% DMSO (100%).

#### 4.6.2. HEK-Slack cells

Viability of HEK-Slack cells was assessed using bright-field microscopy. HEK-Slack cells were cultured in Dulbecco’s modified Eagle’s medium-GlutaMAX with 10% fetal calf serum and 1% penicillin/streptomycin, supplemented with 0.6 mg/mL G-418 (all from Gibco/Thermo Fisher Scientific). Cells were seeded at a density of 1,200 cells/well in 384 well plates in culture medium (cell culture microplate, PS, f-bottom, µClear®, 781091; Greiner). Cell confluence was measured using the IncuCyte® S3 (Satorius) over 24 h. Compounds were tested in biological duplicates. Cell confluence was normalized to cells treated with 0.1% DMSO (100%).

### 4.7. Metabolic stability assay

The solubilized test compound (5 μL, final concentration 10 μM) was preincubated at 37 °C in 432 μL phosphate buffer (0.1 M, pH = 7.4) together with 50 μL NADPH regenerating system (30 mM glucose-6-phosphate, 4 U/mL glucose 6-phosphate dehydrogenase, 10 mM NADP, and 30 mM MgCl_2_). After 5 min, the reaction started upon the addition of 13 μL of microsome mix from the liver of Sprague-Dawley rats (Invitrogen; 20 mg protein/mL in 0.1 M phosphate buffer) in a shaking water bath at 37 °C. The reaction was stopped by adding 500 μL ice-cold methanol at 0, 15, 30, and 60 min. The samples were centrifuged at 5,000 × *g* for 5 min at 4 °C, and the test compound was quantified from the supernatants by HPLC; the composition of the mobile phase was adapted to the test compound in a range of MeOH 40−90% and water (0.1% formic acid) 10−60%. Flow rate:1 mL/min; stationary phase: Purospher STAR, RP18, 5 μm, 125 × 4; precolumn: Purospher STAR, RP18, 5 μm, 4 × 4; detection wavelength:254 and 280 nm; injection volume:50 μL. Control samples were used to evaluate the stability of the test compounds in the reaction mixture. The first control was without NADPH, which is needed for the enzymatic activity of the microsomes, the second control was with inactivated microsomes (incubated for 20 min at 90 °C), and the third control was without the test compound (to determine the baseline). The amounts of test compounds were quantified using an external calibration curve.

### 4.8. Dopamine D_2_/D_3_ receptor affinity assay

CHO cells stably expressing human dopamine D_2short_ (D_2_R) and D_3_ receptor (D_3_R) were washed and harvested using PBS. Cells were centrifuged (3,000 × *g*, 10 min, 4 °C) and homogenized with an Ultraturrax® homogenizer in ice-cold D_2_/D_3_R binding buffer (10 mM MgCl_2_, 10 mM CaCl_2_, 5 mM KCl, 120 mM NaCl, and 50 mM Tris, pH 7.4). The cell membrane homogenate was centrifuged (20,000 × *g*, 20 min, 4 °C) and the resulting cell pellet was resuspended in the binding buffer. Crude membrane extracts were kept at −80 °C until use. Before starting the experiments, cell membranes were thawed, homogenized by sonication (3 × 10 s), and stored in ice-cold binding buffer. For radioligand binding assays, membrane extracts (25 and 20 μg/well in a final volume of 0.2 mL binding buffer for D_2_R and D_3_R, respectively) were incubated with [^3^H]-spiperone (0.2 nM final concentration) and various concentrations of the test ligand for 120 min at room temperature. Assays were performed at least in triplicate with appropriate concentrations ranging from 0.01 nM to 100 μM of the test compound. Non-specific binding was determined using 10 μM haloperidol. By filtering through GF/B filters pretreated with 0.3% (m/v) polyethyleneimine, the bound radioligand was separated from free radioligand using a cell harvester. Radioactivity was measured using liquid scintillation counting. Data were evaluated using the GraphPad Prism 9 software (San Diego, CA, USA) with a nonlinear regression fit.

### 4.9. Histamine H_1_ receptor affinity assay

For membrane preparations of CHO cells stably expressing the human histamine H_1_ receptor (H_1_R), cells were washed and harvested with PBS buffer, homogenized by sonication (3 × 15 s) in ice-cold HEPES-H_1_R binding buffer (20 mM HEPES, 10 mM MgCl_2_, and 100 mM NaCl), followed by a centrifugation step (20,000 × *g*, 30 min, 4 °C). The resulting cell pellet was resuspended in binding buffer and homogenized using a hand potter. Membrane extracts were stored at −80 °C. Prior to use, the membranes were handled as described for D_2_R/D_3_R. For radioligand binding assays, membranes (40 μg/well in a final volume of 0.2 mL binding buffer) were incubated with [^3^H]-pyrilamine (1 nM final concentration) and different concentrations of test ligand for 120 min at room temperature. Assays were performed at least in duplicates with appropriate concentrations between 0.1 nM and 100 μM of the test compound. Nonspecific binding was determined using 10 μM chlorphenamine maleate. The subsequent procedure was performed as described for D_2_R/D_3_R.

### 4.10. Slack expression and purification

The gene encoding isoform 1 of human Slack (*KCNT1*; NCBI ref. seq. NM_020822.3, residues 1–1235) were cloned into the mammalian overexpression vector pcDNA3.1 (Clontech) together with a green fluorescent protein and a twin strep-II-tag sequence for affinity purification at the C-terminus. A TEV protease cleavage sequence was included between the Slack and the GFP sequences to remove the tags. For large-scale expression, HEK293F cells were transiently transfected with the Slack-encoding plasmid using polyethylenimine (PEI) (DNA to PEI ratio of 1:3) and grown in Freestyle™ medium (Thermo Fischer). Seventy-two hours post-transfection, the cells were harvested by centrifugation at 1,000 × *g* for 5 min. Subsequently, the cells were resuspended in ice-cold homogenization buffer (20 mM HEPES, pH 7.6, 350 mM KCl, 10 mM CaCl2, 10 mM MgCl2, and 5 mM DTT) supplemented with 1× complete protease inhibitor mix (Serva), 1% (w/v) DDM, 0.1% (w/v) CHS (Anatrace), and 100 mU benzonase. Solubilization was done for 120 min on an overhead rotator at 4 °C. The lysate was centrifuged at 100,000 × *g* for 1 h and 4 °C to remove the insolubilized membranes and cell debris. The supernatant was then batch-incubated with 250 μL bed volume per L cell culture of Strep-Tactin agarose (IBA Lifesciences) pre-equilibrated with wash buffer (20 mM HEPES, pH 7.6, 350 mM KCl, 10 mM MgCl2, 1× complete protease inhibitor mix, 0.1% (w/v) DDM, 0.01% (w/v) CHS, and 100 µg/mL bovine brain lipids (BBL)) for 4 h at 4 °C. The beads were then collected on a gravity flow column and washed thrice with ten column volumes of ice-cold wash buffer. The remaining protein was digested with TEV protease (20:1 w/w ratio) on the column overnight at 4 °C to remove the affinity tag and then eluted with two column volumes of wash buffer. The eluate was concentrated to 500 μL using an Amicon Ultra-15 (Merck Millipore) and further purified by size-exclusion chromatography on a Superdex 200 10/300 column (GE Life Sciences) in SEC buffer (20 mM HEPES pH 7.6, 350 mM KCl, 0.05% DDM, and 10 μg/mL BBL). Peak fractions were pooled and concentrated to 5 μM (0.7 mg/mL) final concentration using an Amicon Ultra-15 (Merck Millipore) for further analysis.

### 4.11. Differential Scanning Fluorometry (DSF)

The melting temperature was determined from the fluorescence of 7-diethylamino-3-(4-maleimidophenyl)-4-methylcoumarin (CPM) dye. A total of 22.5 μL of 5 µM Slack in SEC buffer was mixed with 22.5 μL of the compounds (compound 6, loxapine, or Slack-inhibiting compound 31 from ref^[27]^) diluted in SEC buffer in the range of 0–200 μM and incubated at 4 °C for 4 h. Five microliters of the CPM dye (15 µg/mL CPM final concentration) in the SEC buffer was added to the protein compound mixture and further incubated for 30 min on ice in the dark. The samples were centrifuged at 15,000 × *g* at 4 °C for 10 min to remove aggregates. Melting curves were recorded in an RT-PCR (Thermo Fischer) using a temperature range of 10–80 °C in 1 °C-increments, increasing every 75 s. The CPM dye was excited at a wavelength of 365 nm and the emission was measured at 470 nm.

### 4.12. In vitro safety panel

A SafetyScreen44 panel was conducted by Eurofins Cerep (Celle l’ Evescault, France; item P270) to identify interactions with 44 preselected pharmacological targets. Loxapine and compound 6 were tested at 10 µM. Measurements were performed in duplicate. The results are expressed as the percent inhibition of control-specific binding obtained in the presence of the test compounds.

### 4.13. Pharmacokinetics

Pharmacokinetic experiments were performed using Bienta (Enamine Biology Services, Kiev, Ukraine). Study design, animal selection, handling, and treatment were performed in accordance with the Enamine PK study protocols and the Institutional Animal Care and Use Guidelines. Male C57BL/6J mice (11–16 weeks old) were used in this study. The animals were fasted for 4 h before dosing. Compounds (dissolved in a solution of 25% 2-hydroxypropyl-β-cyclodextrin in saline) were administered via the i.p., p.o., or i.v. route, and blood and brain tissue were taken at different time points thereafter (0.25, 0.5, 1, 2, and 4 h after i.p. administration; 0.25, 0.5, 1, 2, 4, and 8 h after p.o. administration; 0.083, 0.25, 0.5, 1, 2, and 4 h after i.v. administration). Each timepoint treatment group included three animals, and a control group was employed, including one animal.

Before collecting blood and brain tissue, the mice were anesthetized with 2,2,2-tribromoethanol (150 mg/kg i.p.). In studies with i.p. compound administration, the chest was opened, an incision was made in the right atrium, whole blood was collected into tubes containing K2EDTA, and the animals were perfused with saline (10 mL) prior to the excision of the brain tissue (left lobe). In studies with p.o. or i.v. delivery of compounds, blood was collected from the orbital sinus in microtainers containing K_3_EDTA, the animals were sacrificed by cervical dislocation, and the brain tissue (left lobe) was excised. After collection, the blood samples were centrifuged for 10 min at 3,000 rpm, and the brain samples were weighed. All samples were immediately processed, flash-frozen, and stored at −70 °C until subsequent analysis.

The compound levels in the plasma and brain samples were determined using liquid chromatography–tandem mass spectrometry (LC-MS/MS) by a staff member from Bienta. Pharmacokinetic data analysis was performed using non-compartmental, bolus injection or extravascular input analysis models with WinNonlin 5.2 (PharSight). Data below the lower limit of quantitation were presented as missing data to improve the validity of the half-life calculations. For each treatment condition, the final concentration values obtained at each time point were analyzed for outliers using Grubbs’ test, with the level of significance set at *P* < 0.05. The sum of the AUC_0-inf_ (area under the concentration-time curve from zero to time infinity) was used to estimate the ratio_brain/plasma_ (concentration ratio of the compound in the brain and plasma).

### 4.14. Behavioral tests

#### 4.14.1. Animals

Experiments were performed in 8–16 week-old C57BL/6N mice (Charles River, Sulzfeld, Germany) and Slack^-/-^ mice^[24]^ of either sex. The animals were housed under a 12 h light/dark cycle with access to food and water *ad libitum*. All experiments adhered to the Animal Research: Reporting on In Vivo Experiments (ARRIVE) guidelines and were approved by and conducted in accordance with the regulations of the local Animal Welfare authorities (Regierungspräsidium Darmstadt, Germany; approval number V54-19c20/15-FR/1013 and FR/2011). All behavioral studies were conducted during the light cycle of the day at room temperature (20–24 °C) by an observer blinded for the treatment of the animals and/or their genotype.

#### 4.14.2. Accelerating rotarod and vertical pole test

Mice were placed on a rotarod treadmill (Ugo Basile, Italy) at increasing speed (4–40 rpm over 300 s) and trained for 4–5 consecutive days. Only mice that reached 300 s without falling off during the training sessions were included in the experiment. In the vertical pole test, the mice were placed head-upward on top of a vertical pole with a rough surface (diameter, 1 cm; height, 40 cm), and the time until the animals reached the ground was recorded (cut-off time 20 s). Only mice that reached the ground within 10 s in the baseline measurements were included in the experiment. After baseline measurements without compound administration, compounds or vehicle (10% 2-hydroxypropyl-β-cyclodextrin (ITW Reagents) in 0.9% saline; used for compound 6, loxapine, and morphine in all behavioral tests in this study) were i.p. administered, and 15 min later, the accelerating rotarod followed by the vertical pole test were performed. The means of three trials at each time point were calculated for further analyses.

#### 4.14.3. Acute itch behavior

Three to four days before the day of the experiment, the fur on the dorsolateral aspect of the neck was shaved under brief isoflurane anesthesia. On the testing day, the mice were habituated to a Plexiglas cylinder (30 cm in diameter) for 30 min. Compound 6 or vehicle were i.p. or p.o. administered, and 15 min later, the pruritogens chloroquine (200 μg), SLIGRL (100 µg), β-alanine (100 µg), or histamine (800 μg), all dissolved in 0.9% saline (20 μL), were injected subcutaneously into the nape of the neck.^[29]^ Numbers of scratching bouts directed towards the nape of the neck was assessed over 30 min by videotaping. In experiments with the Slack inhibitor, compound 31 (ref^[27]^; 30 mg/kg in 0.9% NaCl with 2% DMSO and 10% Kolliphor) was i.p. administered 5 min prior to compound 6.

#### 4.14.4. Allergic contact dermatitis model

To induce persistent itch, a model of allergic contact dermatitis^[32]^ was used. The fur on the dorsolateral aspect of the neck was shaved under brief isoflurane anesthesia. Three to four days later, the shaved skin area was painted with 100 μL of 0.15% 2,4-dinitrofluorobenzene (DNFB) in acetone/olive oil (3:1). After 10–11 days, the skin was shaved again, and the DNFB solution was painted 3–4 days later (i.e., two painting sessions 14 days apart). Ninety minutes after the second painting, the mice were habituated to a plexiglass cylinder (30 cm diameter) for 15 min. Compound 6 or vehicle was administered i.p., and 15 min later, spontaneous scratching was recorded over 30 min by videotaping.

#### 4.14.5. Atopic dermatitis model

Chronic itching was induced using the MC903 model.^[47]^ The fur on the dorsolateral aspect of the neck was shaved under brief isoflurane anesthesia. Five days later, 20 μL MC903 (0.2 mM; Calcipotriol, Tocris) solved in absolute ethanol was applied to the skin at the back of the neck under brief anesthesia for 7 consecutive days. On day 8, the mice were habituated for 30 min in a plexiglass cylinder (30 cm in diameter). Compound 6 or vehicle was administered i.p., and 15 min later, spontaneous scratching was recorded over 30 min by videotaping.

#### 4.14.6. Pulse oximetry

To measure cardiovascular function, a pulse oximeter (MouseOX Plus, Starr Life Sciences Corp.) was used in conscious, freely moving mice. After three consecutive days of habituating the mice to the collar clip for at least 30 min, baseline measurements were performed. Compounds or vehicle were i.p. administered, and after a resting time of 5 min, the heart rate, respiratory rate, and arterial O_2_ saturation were recorded over 30 min. All the datasets were sampled at 1 Hz, error-corrected according to the manufacturer’s instructions, and averaged over 300 s for each time point.

#### 4.14.7. Hargreaves test

Mice were placed in plastic chambers on a heated glass panel (32 °C) of a Plantar Test device (Hargreaves method; IITC Life Science, CA) and acclimated for 1 h. An infrared heat source (intensity set to 25) was applied to the plantar surface of the hind paw, and the time until the animal elicited a withdrawal response was determined. Each hind paw was measured 4–5 times with a minimum interval of 20 s between measurements, and the mean value was calculated. Measurements that reached the cutoff time (20 s) were excluded. Thermal sensitivity was assessed at baseline and 15 min after i.p. administration of compound 6 or vehicle.

#### 4.14.8. Von Frey filament test

The mice were placed in plastic chambers on a wire mesh grid and acclimated for 1 h. Mechanical sensitivity was assessed using von Frey filaments (Ugo Basile) with increasing forces (0.04, 0.07, 0.16, 0.4, 0.6, 1, 1.4, 2, and 4 g) applied to the plantar surface of the hind paw. A withdrawal response within 1 s of a slight hair bending was recorded as a positive reaction. Each filament was applied 10 times within 10 min and both hind paws were measured equally. The total number of responses per force was recorded as the percentage of withdrawals per mouse. After baseline measurement, compound 6 or vehicle was administered intraperitoneally, and 15 min later, a second measurement was performed.

### 4.15. Statistical Analysis

Statistical analysis was performed using Prism 9 (GraphPad). No statistical methods were used to pre-determine the sample sizes; however, our sample sizes were similar to those reported in previous publications^[29, 31, 32, 34]^ and standard practices in the field. Kolmogorov-Smirnov or Shapiro-Wilk tests were used to assess the normal distribution of data within groups. Normally distributed data were analyzed using two-tailed unpaired or paired t-test, one-way-ANOVA, or two-way MC ANOVA and are expressed as the mean ± standard error of the mean (SEM) or single data points with the mean ± SEM. Nonparametric data were analyzed using a two-tailed Mann-Whitney test or Kruskal-Wallis test and presented as single data points with medians and interquartile ranges. Outliers were identified using the ROUT method (Q = 1%), and one outlier was excluded from the analysis (APD90 recordings in Figure 5D) in the entire study. For all statistical tests, a probability value *P* < 0.05 was considered statistically significant. Asterisks in the figures indicate: \**P* < 0.05, \*\**P* < 0.01, and \*\*\**P* < 0.001. Data from FluxOR experiments (Figure 1A) were analyzed applying a standard, logistic, nonlinear regression implemented in Prism 9 to get dose-response curves representing EC_50_ and E_MAX_ values ± 95% CI. In the *in vivo* experiments, behavioral tests were performed by observers blinded to the treatment groups or genotypes. The treatment groups were randomized and evenly distributed across cages and sexes. Details of the analyses, including the statistical test, post-hoc test, number of replicates for each analysis, and number of animals per group, are indicated in the figure legends.

## Acknowledgements

We thank Astrid Kaiser, Sylvia Oßwald and Cyntia Schäfer for excellent technical assistance.

## Author Contributions

A.B. performed FluxOR assays, electrophysiological recordings, *in vivo* experiments, analyzed data, prepared the figures and assisted with preparing the manuscript. W.F.Z. designed and synthesized compounds and assisted with preparing the manuscript. C.F. established the FluxOR assay, performed initial FluxOR and *in vivo* experiments, and analyzed these data. V.H.-O. designed and synthesized compounds. J.H. established and supervised the FluxOR assay. S.S and I.H. performed and/or evaluated DSF analyses. M.D., A.F. and H.S. performed dopamine and histamine binding assays. A.M., S.K. and S.M. conducted and/or evaluated cytotoxicity measurements. M.H. assisted in electrophysiological recordings. K.M. supervised the electrophysiological recordings. R. Lu performed *in vivo* experiments. R. Lukowski and P.R. provided Slack knockout mice and analytical tools. D.S. contributed to the conceptualization of the project and edited the manuscript. H.S. conceived the project. E.P. conceived the project, supervised compound design and synthesis, analyzed data and wrote the manuscript. A.S. conceived the project, designed und supervised the *in vivo* experiments, analyzed data and wrote the manuscript. All authors revised and approved the manuscript.

---

**Figure S1.**
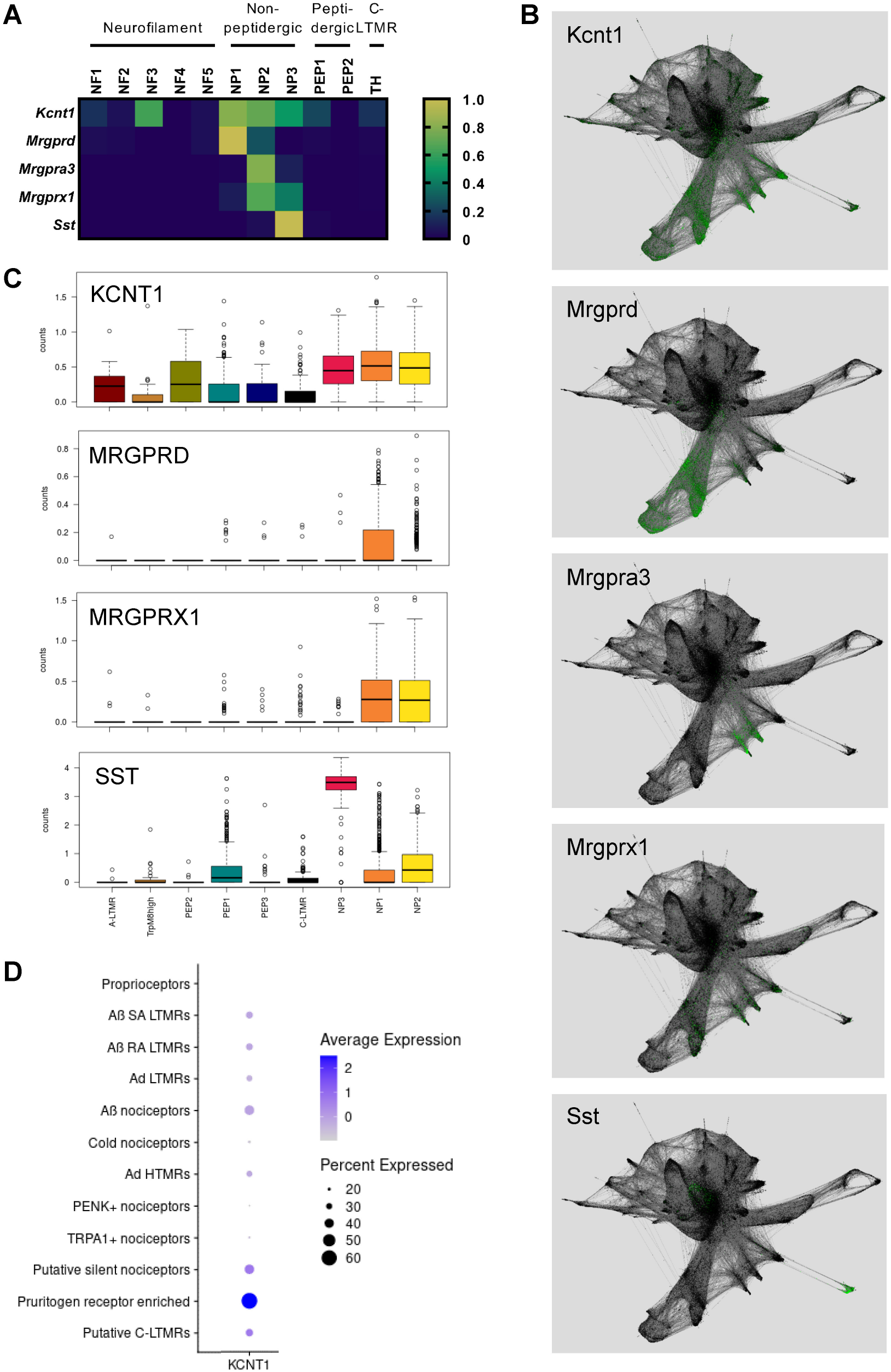
Expression of Kcnt1 and critical itch receptors across sensory neuron subsets. Presented are data from published single-cell RNA-seq studies. A) Expression pattern in mouse DRG neurons (6-8 week old).^[1]^ B) Expression pattern in mouse DRG neurons (embryonic day 11.5 to postnatal day 42) presented as a force-directed layout.^[2]^ Downloaded from: https://kleintools.hms.harvard.edu/tools/springViewer_1_6_dev.html?datasets/Sharma2019/all. C) Expression pattern in non-human primate DRG neurons (5-14 year old).^[3]^ Downloaded from: https://ernforsgroup.shinyapps.io/macaquedr. D) Expression pattern in human DRG neurons (24-65 year old).^[4]^ Downloaded from: https://sensoryomics.shinyapps.io/RNA-Data/.

**Figure S2.**
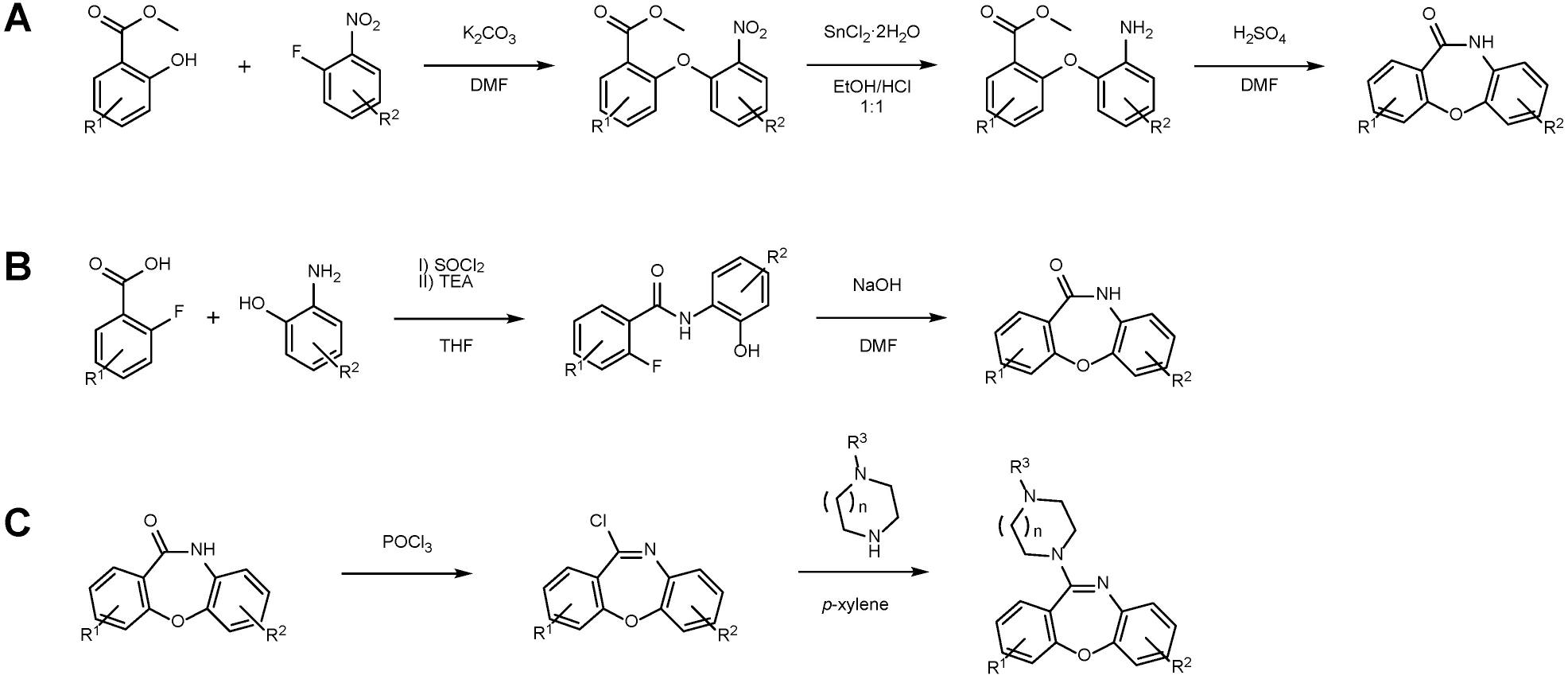
Chemical synthesis scheme of new compounds. A) General synthesis of lactam starting from methyl esters of salicylic acid and 2-fluoronitrobenzene derivatives. B) General synthesis of lactam starting from 2-fluorobenzoic acids and 2-aminophenol derivatives. C) General synthesis of loxapine derivatives starting from lactam.

**Figure S3.**
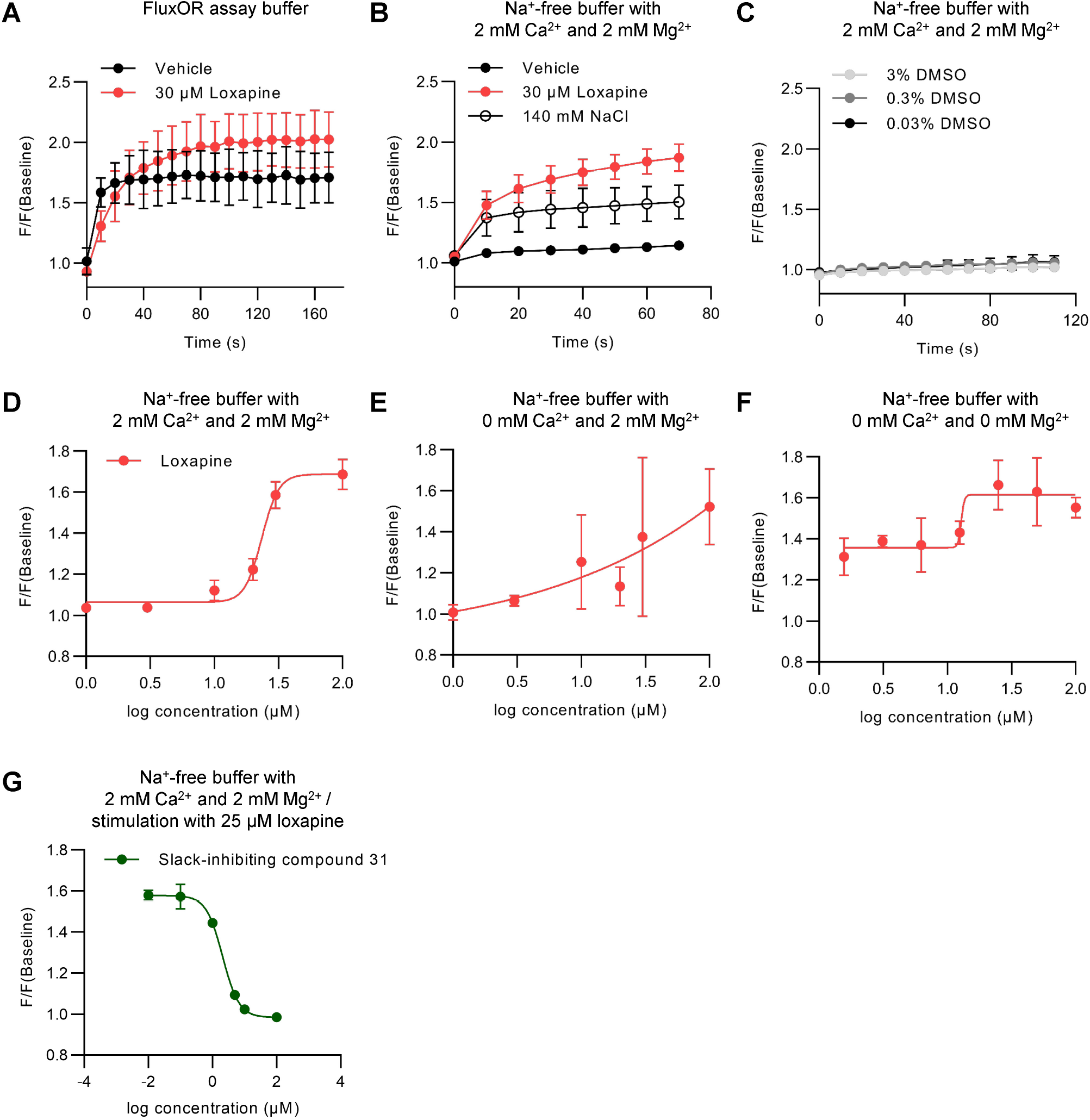
Establishing a modified version of the FluxOR assay. Cultured HEK293 cells stably expressing human Slack (HEK-Slack cells) were incubated with compounds in different buffers. A) In the buffer provided with the FluxOR assay kit, Slack activation (indicated as increased F/F(Baseline) ratio) was detected after incubation with both loxapine and vehicle (FluxOR assay buffer containing 0.03% DMSO). As Slack is activated by Na^+^, the vehicle-induced Slack activation was most likely mediated by Na^+^ present in the FluxOR assay buffer and in the vehicle. B) In a Na^+^-free buffer (with replacement of NaCl by choline chloride), Slack activation was observed after incubation with 30 µM loxapine and 140 mM NaCl, but not after incubation with vehicle (Na^+^-free buffer with 2 mM Ca^2+^, 2 mM Mg^2+^ and 0.03% DMSO). C) DMSO at a concentration of 0.03-3% did not activate Slack in a Na^+^-free buffer with 2 mM Ca^2+^ and 2 mM Mg^2+^. D) Concentration-response experiments with loxapine in a Na^+^-free buffer with 2 mM Ca^2+^, 2 mM Mg^2+^ and 0.03% DMSO yielded an EC_50_ value of 23.45 µM in the first preliminary experiments. E,F) As Slack is inhibited by bivalent cations, further control experiments with loxapine in a Na^+^ and Ca^2+^ free buffer (E) and in a Na^+^, Ca^2+^ and Mg^2+^ free buffer (F) containing 0.03% DMSO were conducted, which however did not result in appropriate concentration-response curves. G) Concentration-response experiments with a Slack inhibitor (compound 31 from ref ^[5]^) in a Na^+^-free buffer with 2 mM Ca^2+^, 2 mM Mg^2+^ and 0.03% DMSO and pre-stimulation with 25 µM loxapine revealed that compound 31 inhibited the F/F(Baseline) ratio in a concentration-dependent manner (IC_50_ = 2.1 µM), confirming that the readout depends on Slack. Therefore, all further experiments with the FluxOR assay in this study, which are presented in Figure 1A, were performed using a Na^+^-free buffer with 2 mM Ca^2+^, 2 mM Mg^2+^ and 0.03% DMSO. All conditions were measured at least in duplicate and data are shown as mean ± SD.

**Figure S4.**
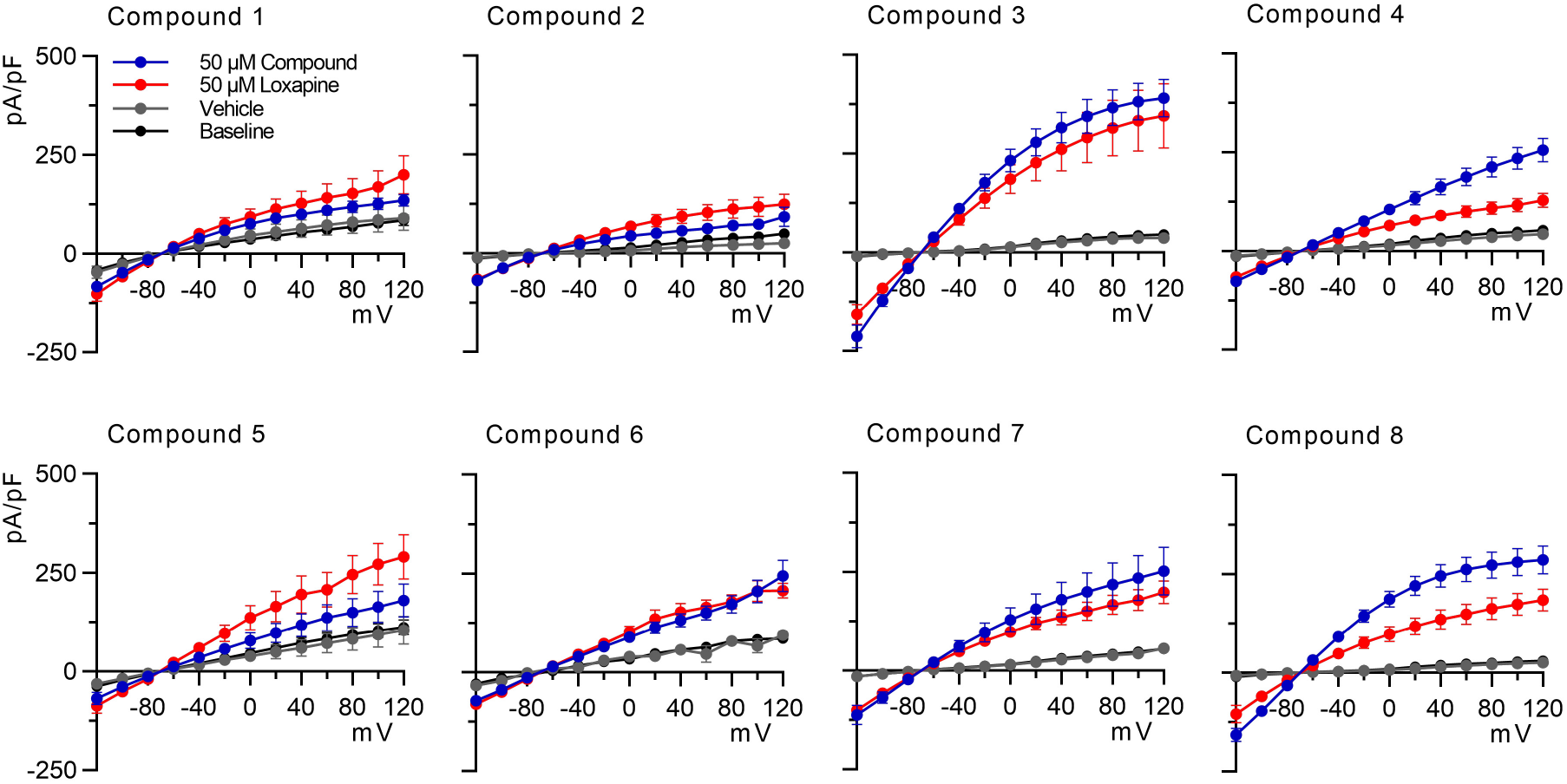
New compounds evoke Slack-mediated potassium currents. Current-voltage (I-V) curves from patch-clamp experiments that are presented in Figure 1B. Whole-cell voltage recordings on HEK-Slack cells were performed at baseline and after incubation with a new compound (50 µM), loxapine (50 µM) or vehicle (external solution containing 0.03% DMSO). Loxapine was incubated in each series of experiments as a positive control. *n* = 5-24 cells per group. Data are shown as mean ± SEM.

**Figure S5.**
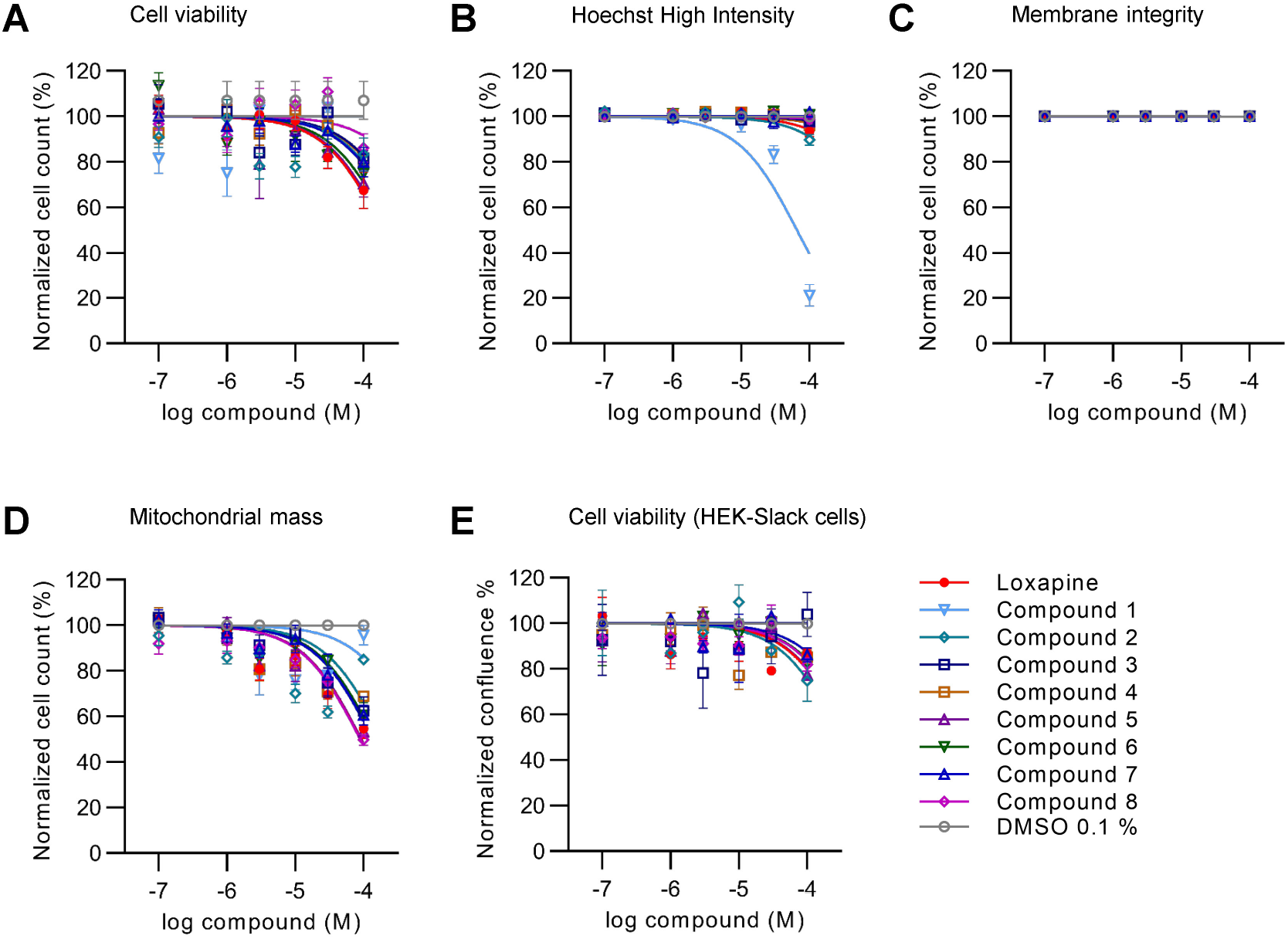
Cytotoxicity of new compounds. A–D) A live-cell phenotypic screening assay that allows for simultaneous investigation of cellular viability, nuclear morphology, mitochondrial mass and membrane integrity^[6]^ was performed in HEK293 cells after 6 h incubation of new compounds at different concentrations. Cell counts were normalized against cells treated with 0.1 % DMSO (100%). A) None of the compounds decreased the cell viability by more than 50%, which is deemed a relevant impairment.^[6]^ B) A decrease of normalized cell count to more than 50% based on Hoechst High Intensity objects, which indicates compound precipitation,^[7]^ was only detected for compound 1 at 100 µM. C) No alterations in membrane integrity were detected. D) A decrease in mitochondrial mass by more than 50%, indicating cellular stress,^[6]^ was only observed for compound 8 at 100 µM. E) In HEK-Slack cells, none of the compounds significantly impaired the confluence after 24 h incubation. Data are shown as mean ± SEM of biological duplicates.

**Figure S6.**
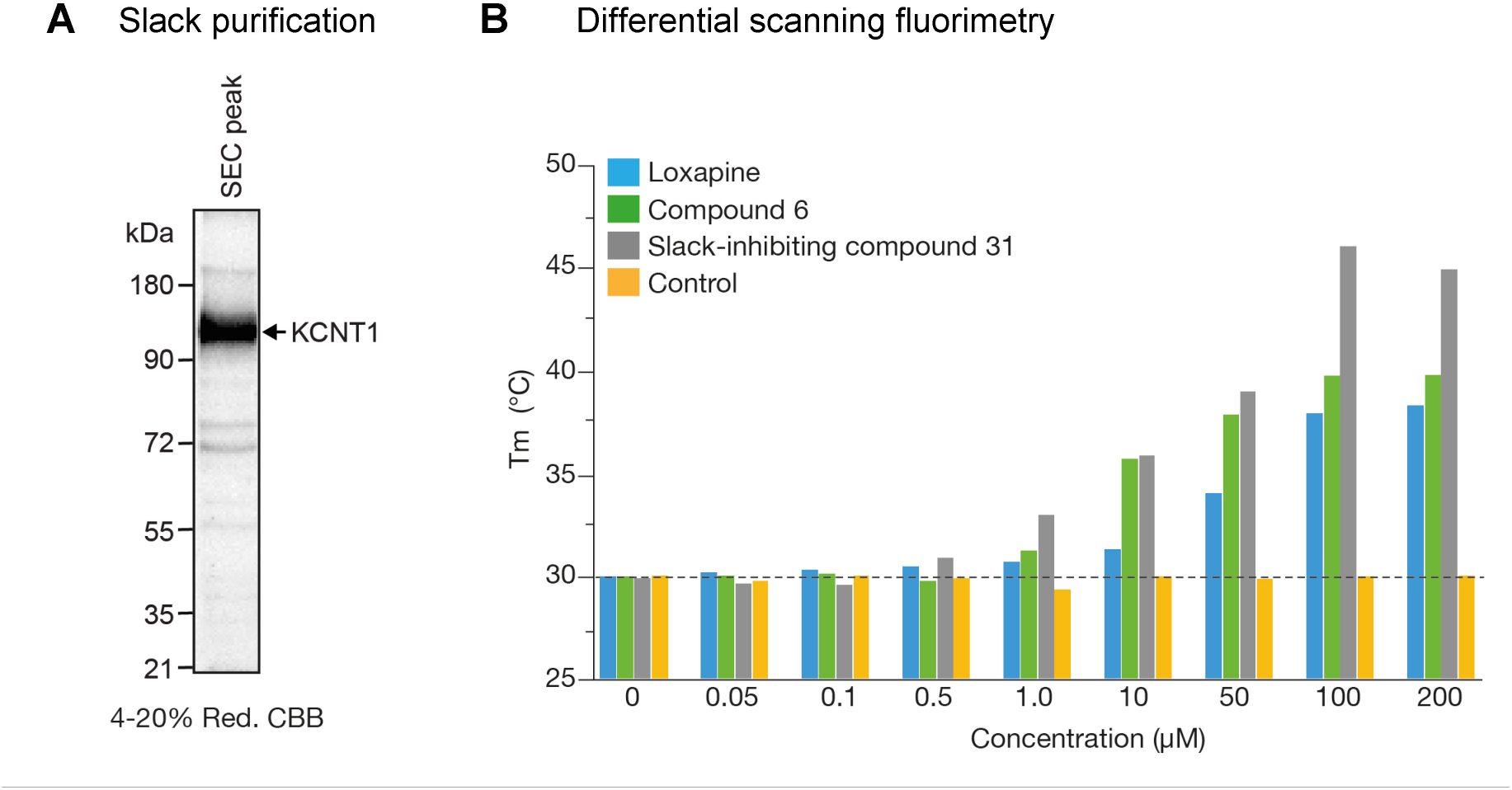
Target binding of compound 6 and loxapine. A) Slack was purified from HEK293F cells transiently expressing Slack via affinity purification and size-exclusion chromatography. B) Melting temperatures of purified Slack incubated with an increasing concentration of compound 6, loxapine and a Slack inhibitor (compound 31 from ref^[5]^) that was used as positive control. Purified Slack in buffer without any compound was used as a negative control. The melting temperature (*T_m_* in °C) was determined with differential scanning fluorimetry (DSF) and plotted against the concentration of the compounds used. The dashed line indicates the *T_m_* of purified Slack without compound.

**Figure S7.**
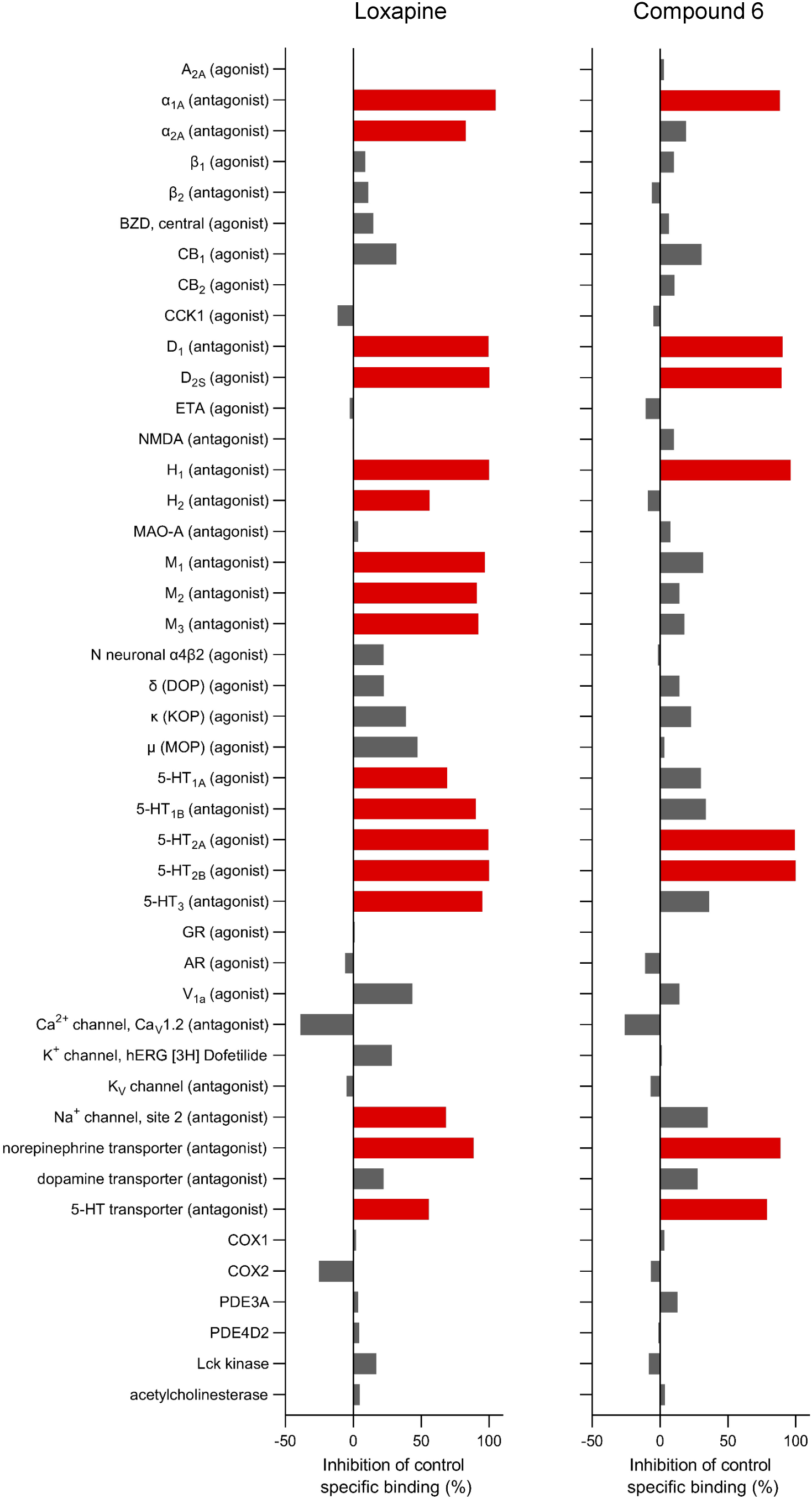
*In vitro* pharmacology screening of compound 6 and loxapine. Compounds were tested at a concentration of 10 µM in binding and enzyme and uptake assays of 44 targets (mostly human; except BZD, NMDA, MAO-A, Ca^2+^ channel, K_V_ channel, and Na^+^ channel, which were obtained from rat specimens). Compound binding was calculated as a % inhibition of the binding of a radioactively labeled ligand (agonist or antagonist, as indicated in brackets) specific for each target. Compound enzyme inhibition effect was calculated as % inhibition of control enzyme activity. Results showing an inhibition (or stimulation for assays run in basal conditions) higher than 50% are considered to represent significant effects of the test compounds and are presented in red. Results showing an inhibition or stimulation between 25% and 50% (indicative of weak to moderate effects) and those lower than 25% (considered mostly attributable to variability of the signal around the control level) are presented in grey. Measurements were performed in duplicate.

**Figure S8.**
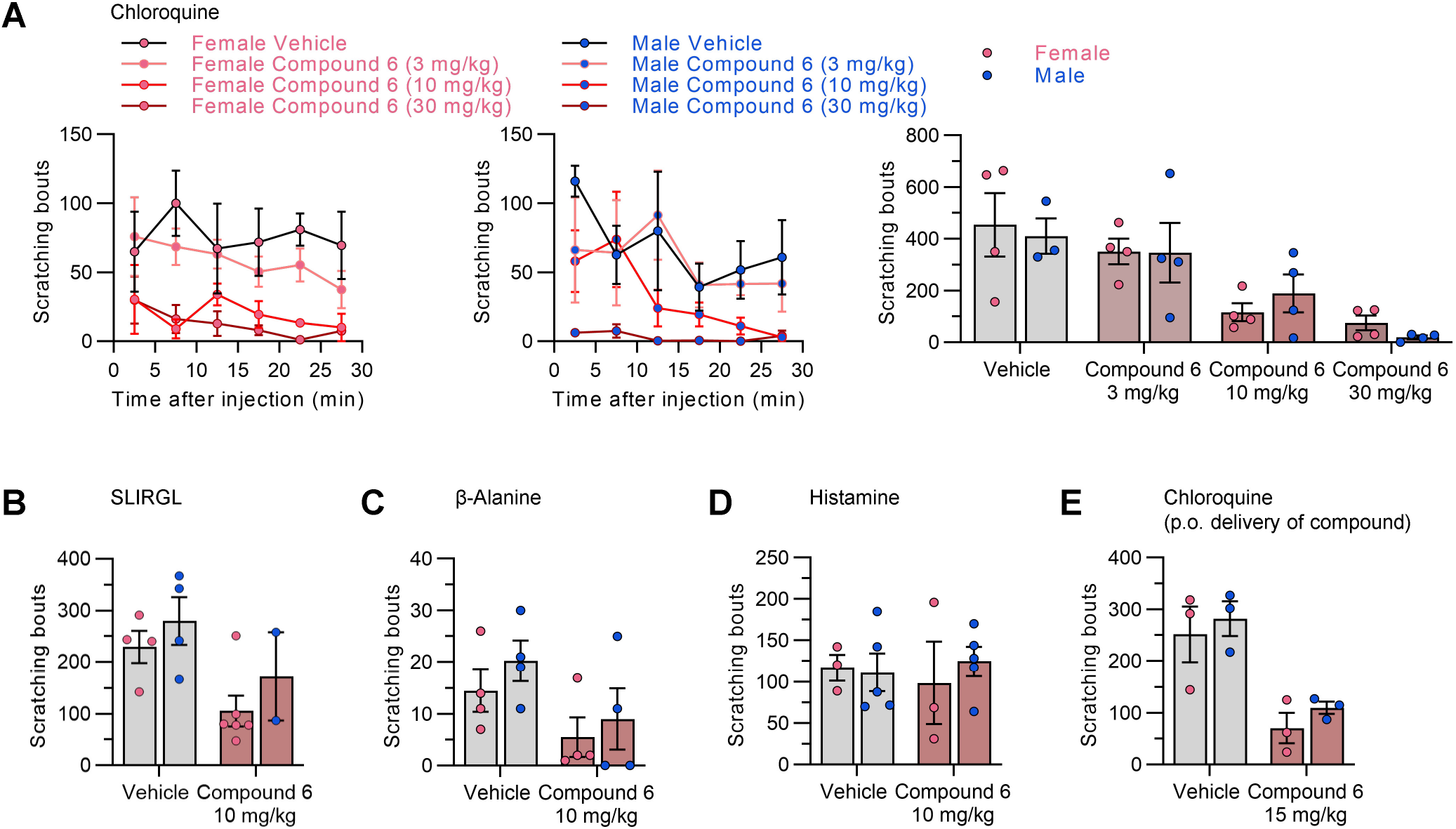
Sex-related effects of compound 6 in acute itch models. Breakdown of results in male and female mice from (A) Figure 3C, (B) Figure 3E, (C) Figure 3F, (D) Figure 3G, and (E) Figure 3H. No obvious sex-related differences were observed in any assay. Statistical significance was assessed by one-way-ANOVA with Dunnett’s correction. Data represent the mean ± SEM (*n* = 2–5).

**Figure S9.**
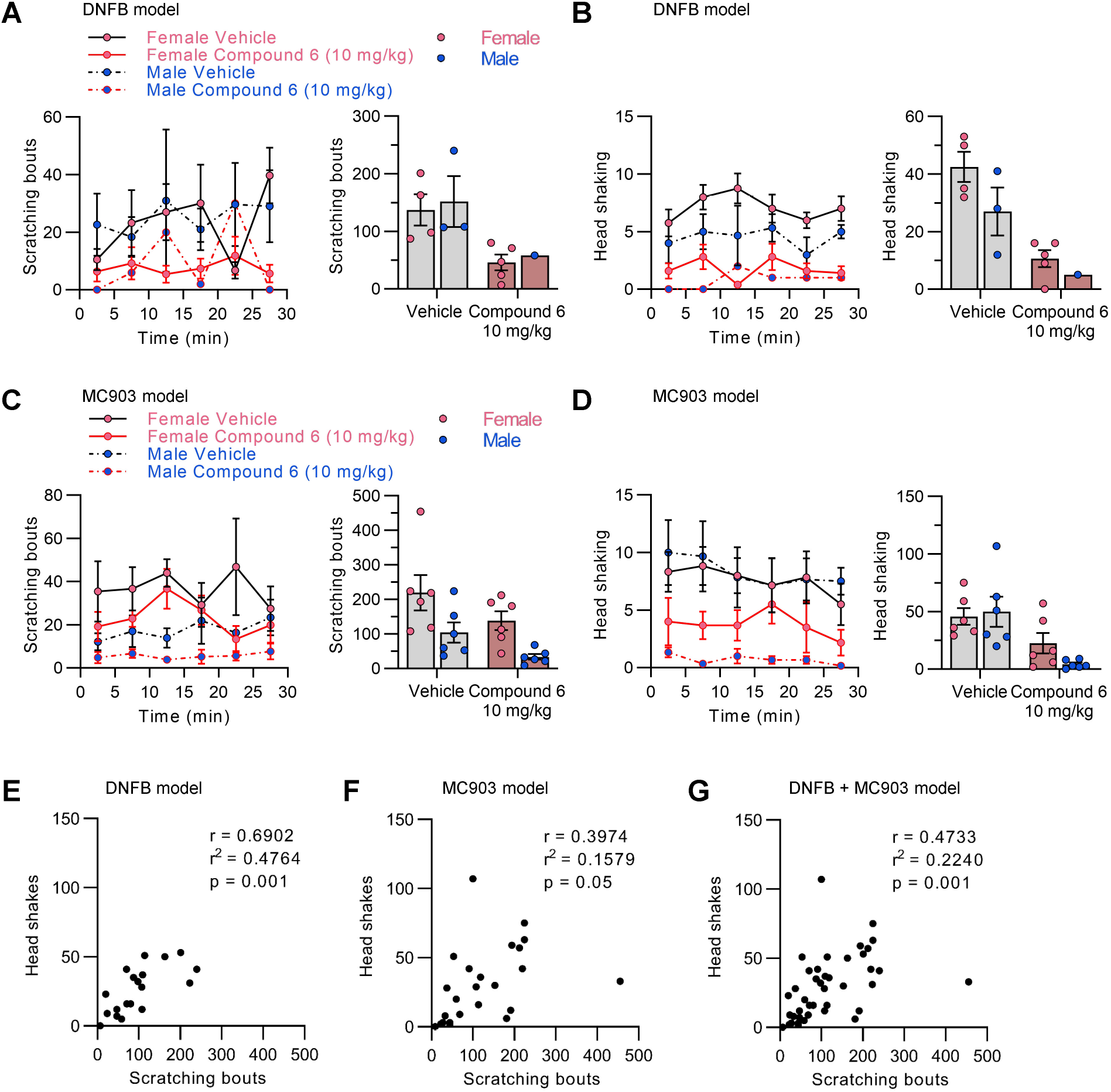
Sex-related effects of compound 6 and correlation of behavioral outcomes in chronic itch models. A-D) Breakdown of results in male and female mice from (A) Figure 4B, (B) Figure 4C, (C) Figure 4E, and (D) Figure 4F. There was a tendency toward reduced chronic itch behavior in males compared to females (head shaking in the DNFB model and scratching in the MC903 model), albeit not significant. Statistical significance was assessed by one-way-ANOVA with Dunnett’s correction. Data represent the mean ± SEM (*n* = 1–6). E–G) Correlation of number of head shakes with number of scratching bouts (E) in the DNFB model (Figure 4B,C), (F) in the MC903 model (Figure 4E,F) and (G) in both models. Statistical significance was assessed by a Pearson correlation.

**Figure S10.**
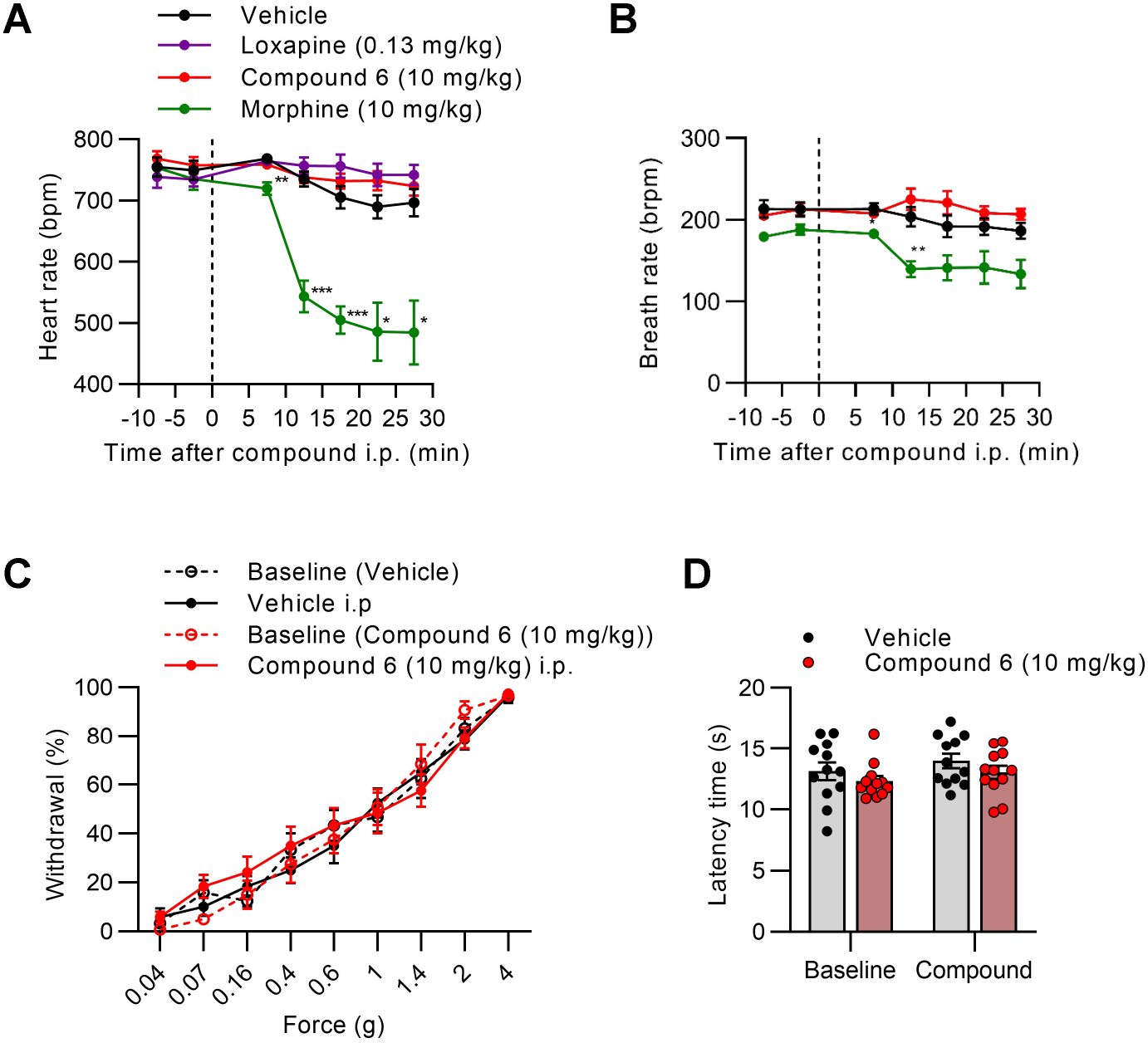
Potential side effects of compound 6. A,B) Pulse oximetry on non-anesthetized mice using a MouseOX Plus device revealed that i.p. delivery of compound 6 or vehicle did not alter (A) the heart rate or (B) the breath rate, whereas morphine significantly lowered both parameters (*n* = 6; \**P* < 0.05, \*\**P* < 0.01, \*\*\**P* < 0.001 vs vehicle, two-way MC ANOVA and Dunnett test). C,D) Sensing of (C) mechanical stimuli (von Frey filament test) and (D) heat stimuli (Hargreaves test) was not affected after i.p. delivery of compound 6 or vehicle (*n* = 12). Data represent the mean ± SEM.

**Table S1.**
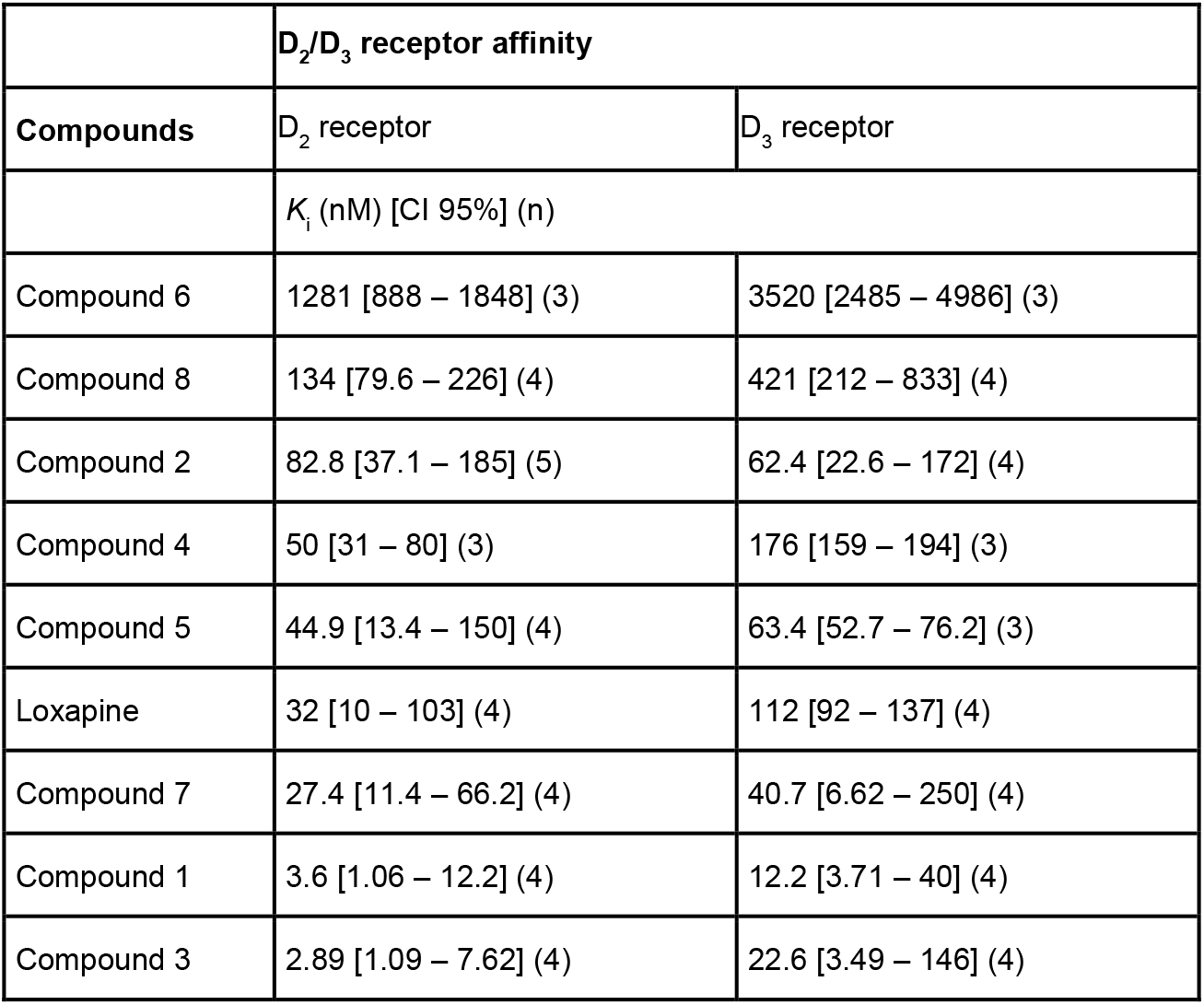
Dopamine D_2_/D_3_ receptor affinity of the new compounds. D_2_/D_3_ receptor affinity values are from data presented in Figure 2B. [^3^H]-Spiperone competition binding assays were performed using a cell membrane preparation of CHO cells stably expressing the human D_2s_ and D_3_ receptor. K_i_ data represent mean with the 95% confidence interval (CI 95%) of *n* independent experiments, each performed in triplicate using seven appropriate concentrations of test compound. Haloperidol was used as a reference compound and to determine non-specific binding at 10 μM.

|  | **D<sub>2</sub>/D<sub>3</sub> receptor affinity** |  |
| --- | --- | --- |
| **Compounds** | D<sub>2</sub> receptor | D<sub>3</sub> receptor |
|  | *K<sub>i</sub>* (nM) [CI 95%] (n) |  |
| Compound 6 | 1281 [888 – 1848] (3) | 3520 [2485 – 4986] (3) |
| Compound 8 | 134 [79.6 – 226] (4) | 421 [212 – 833] (4) |
| Compound 2 | 82.8 [37.1 – 185] (5) | 62.4 [22.6 – 172] (4) |
| Compound 4 | 50 [31 – 80] (3) | 176 [159 – 194] (3) |
| Compound 5 | 44.9 [13.4 – 150] (4) | 63.4 [52.7 – 76.2] (3) |
| Loxapine | 32 [10 – 103] (4) | 112 [92 – 137] (4) |
| Compound 7 | 27.4 [11.4 – 66.2] (4) | 40.7 [6.62 – 250] (4) |
| Compound 1 | 3.6 [1.06 – 12.2] (4) | 12.2 [3.71 – 40] (4) |
| Compound 3 | 2.89 [1.09 – 7.62] (4) | 22.6 [3.49 – 146] (4) |

**Table S2.** Pharmacokinetic parameters of compound 6 and loxapine in mice. Animals were i.p., p.o. or i.v. injected with the compound at the indicated dose and plasma or brain concentration at different time points (0.25, 0.5, 1, 2 and 4 h after i.p. delivery; 0.25, 0.5, 1, 2, 4 and 8 h after p.o. delivery; 0.083, 0.25, 0.5, 1, 2 and 4 h after i.v. delivery) were measured by LC-MS analysis (*n* = 3 mice per group). C_max_: maximum concentration reached in plasma or brain; t_max_: time to reach C_max_; AUC_0-inf_: area under the concentration-time curve from zero to time infinity; t_1/2_: half-life.

| Compound | Route (dose) | PK parameters | Plasma | Brain |
| --- | --- | --- | --- | --- |
| Compound 6 | I.P.<br>(10 mg/kg) | C<sub>max</sub> (ng/mL) | 6280 | 5490 |
|  |  | t<sub>max</sub> (min) | 15 | 15 |
|  |  | AUC<sub>0-inf</sub> (ng*min/mL) | 336000 | 407000 |
|  |  | t<sub>1/2</sub> (min) | 35.0 | 39.7 |
|  | P.O.<br>(10 mg/kg) | C<sub>max</sub> (ng/mL) | 3140 | 2460 |
|  |  | t<sub>max</sub> (min) | 15 | 15 |
|  |  | AUC<sub>0-inf</sub> (ng*min/mL) | 244000 | 225000 |
|  |  | t<sub>1/2</sub> (min) | 67.4 | 71.9 |
| Compound 6 | I.V.<br>(1 mg/kg) | C<sub>max</sub> (ng/mL) | 562 | 493 |
|  |  | t<sub>max</sub> (min) | - | 15 |
|  |  | AUC<sub>0-inf</sub> (ng*min/mL) | 28400 | 31200 |
|  |  | t<sub>1/2</sub> (min) | 52.5 | 62.1 |
| Loxapine | I.P.<br>(2.5 mg/kg) | C<sub>max</sub> (ng/mL) | 143 | 493 |
|  |  | t<sub>max</sub> (min) | 15 | 15 |
|  |  | AUC<sub>0-inf</sub> (ng*min/mL) | 10700 | 54800 |
|  |  | t<sub>1/2</sub> (min) | 61.9 | 82.2 |

**Table S3.** Histamine H_1_ receptor affinity of compound 6 and loxapine. [^3^H]-Pyrilamine competition binding assays were performed using cell membrane preparation of CHO cells stably expressing the human H_1_ receptor. K_i_ data represent mean with the 95% confidence interval (CI 95%) of *n* independent experiments, each performed in triplicate using seven appropriate concentrations of test compound. Chlorphenamine maleate was used as reference compound. Non-specific binding was determined in the presence of 10 µM chlorphenamine maleate.

| Compound | $K_i$ (nM) [CI 95%] (n) |
| --- | --- |
| Chlorphenamine maleate | 9.07 [1.98 – 41.7] (5) |
| Loxapine | 2.72 [0.65 – 11.5] (5) |
| Compound 6 | 575 [137 – 2422] (5) |

## Supplemental protocols for synthesis of compounds

**(A) General synthesis of lactam starting from methyl esters of salicylic acid and 2-fluoronitrobenzene derivatives.**

### GP1: Formation of the diarylether

Potassium carbonate (1.5 equiv) was added to a solution of the corresponding methyl salicylate (1.5 equiv) and the corresponding 2-fluoronitrobenzene (1.0 equiv) in DMF (1.5 M regarding 2-fluoronitrobenzene). The resulting solution was heated in an oil bath to 120 °C overnight. Solvents were evaporated under reduced pressure and the residue taken into water and extracted three times with ethyl acetate. The combined organic phases were dried over magnesium sulfate, filtered, and evaporated. The resulting crude product was purified by flash chromatography.

### GP2: Reduction of the nitro group

A solution of SnCl_2_·2H_2_O (4.0 equiv) in conc. HCl (3 M) was added to a solution of the corresponding methyl 2-(2-nitrophenoxy)benzoate derivative (1.0 equiv) in a mixture of ethanol/conc. HCl 1:1 (0.5 M). The resulting solution was stirred at rt overnight. After that time, the temperature was set to 0 °C and the pH of the reaction solution made slightly basic by addition of sodium carbonate. The resulting solution was then extracted three times with ethyl acetate. The combined organic phases were dried over magnesium sulfate, filtered, and evaporated. The resulting crude product was purified by flash chromatography.

### GP3: Formation of the lactam ring via intramolecular condensation

A solution of the corresponding methyl 2-(2-aminophenoxy) benzoate derivative (1.0 equiv) in DMF (0.2 M) was treated with concentrated sulfuric acid (1.3 equiv) and heated in an oil bath at 120 °C overnight. After that time, the reaction was cooled to 0 °C and a few millilitres of water were added. The precipitated product was then filtered off and dried under vacuum to obtain the crude product, which was used in the next step without further purification.

**(B) General synthesis of lactam starting from 2-fluorobenzoic acids and 2-aminophenol derivatives.**

### GP4: Formation of the amide

To a solution of the corresponding 2-fluorobenzoic acid (1.0 equiv) in THF (1 M) freshly distilled thionylchloride (2.0 equiv) was added and heated to reflux in an oil bath for 2 h. Thereafter, excess thionylchloride and THF were removed under reduced pressure and the residue obtained was taken up in THF (2.5 M) again. This solution was added dropwise to a solution of the corresponding 2-aminophenol derivative (1.0 equiv) and triethylamine (2 equiv) in THF (2.5 M referred to 2-aminophenol derivative) at 0 °C and the reaction mixture was stirred overnight at rt. After that time, the reaction mixture was concentrated under reduced pressure and the residue taken up in ethyl acetate. The organic phase was first washed with an aq HCl solution (2 M), water and saturated aq NaCl solution, then dried over magnesium sulfate, filtered, and evaporated. The resulting crude product was purified by flash chromatography.

### GP5: Formation of the lactam ring via intramolecular nucleophilic aromatic substitution

A solution of the corresponding 2-fluoro-*N*-(2-hydroxyphenyl)benzamide derivative (1.0 equiv) in DMF (0.25 M) was treated with freshly powdered NaOH (1.0 equiv) and heated in an oil bath at 150 °C for 5 h. After that time, the reaction was cooled to 0 °C and a few millilitres of water were added. The precipitated product was then filtered off and dried under vacuum to obtain the crude product, which was used in the next step without further purification.

**(C) General synthesis of loxapine derivatives starting from lactam**

### GP6: Reaction with phosphorus oxychloride

The corresponding lactam (1.0 equiv) was dissolved in freshly distilled phosphorus oxychloride (0.5 M) and *N*,*N*-dimethylaniline (0.6 equiv) was added. The resulting mixture was heated to reflux in an oil bath for 5 h. Thereafter, excess phosphorus oxychloride was removed under reduced pressure and the residue obtained was taken up in toluene and washed once with cold water. The organic phase was dried over magnesium sulfate, filtered, and evaporated to give the crude product, which was immediately used in the next step without further purification.

### GP7: Formation of loxapine derivatives

The corresponding amine (2.0 equiv) was added to a solution of the appropriate imidoylchloride (1.0 equiv) in *p*-xylene (0.15 M regarding imidoylchloride). The resulting mixture was heated in an oil bath at 140 °C for 5 h. Solvents were evaporated, and the crude substance purified by preparative HPLC or flash chromatography. If unsubstituted piperazine or homopiperazine was employed, then alkylation of these derivatives was obtained by GP8

### GP8: Alkylation of loxapine derivatives

A mixture of loxapine derivative (1 equiv), corresponding alkyl chloride (2 equiv) and triethylamine (10 equiv) in acetonitrile (0.15 M) was heated to reflux in an oil bath for 16 h. After completion, the reaction mixture was evaporated, and the residue purified by preparative HPLC or flash chromatography.

## Characterization of compounds

### Methyl 2-methoxy-5-(trifluoromethyl)benzoate (9)

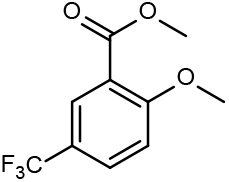

A solution of 2-methoxy-5-(trifluoromethyl)benzoic acid (1.00 g, 4.41 mmol) in MeOH (10 mL) was treated with concentrated sulfuric acid (235 µL, 4.41 mmol) and heated to reflux in an oil bath overnight. The reaction mixture was concentrated under reduced pressure and the residue was taken up in water. The aqueous phase was extracted three times with ethyl acetate and the combined organic phases were dried over magnesium sulfate and filtered. Concentrating the reaction mixture under reduced pressure yielded the title compound as a colourless oil (1.02 g, quantitative). ^1^H NMR (250 MHz, acetone-*d*_6_) δ 7.99 (d, J = 2.4 Hz, 1H), 7.86 (dd, J = 8.8, 2.4 Hz, 1H), 7.36 (d, J = 8.8 Hz, 1H), 3.98 (s, 3H), 3.86 (s, 3H).

### Methyl 2-hydroxy-5-(trifluoromethyl)benzoate (10)

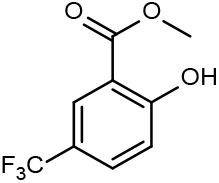

A solution of boron tribromide (7.8 mL, 7.80 mmol) in DCM (1 M) was added to a solution of **9** (939 mg, 3.89 mmol) in DCM (30 mL) at −78 °C and the resulting mixture was first stirred at −78 °C for 5 min, then at 0 °C for 10 min. A few mL of MeOH were added before the reaction mixture was concentrated under reduced pressure and the residue was diluted with ethyl acetate. The organic phase was washed twice with water and once with a saturated aq NaCl solution, dried over magnesium sulfate, and filtered. Concentrating the reaction mixture under reduced pressure yielded the title compound as a brown oil (364 mg, 43%). ^1^H NMR (250 MHz, acetone-*d*_6_) δ 11.11 (s, 1H), 8.15 (d, J = 1.9 Hz, 1H), 7.86 (dd, J = 8.8, 2.4 Hz, 1H), 7.18 (d, J = 8.8 Hz, 1H), 4.03 (s, 3H).

### Methyl 5-chloro-2-(2-nitrophenoxy)benzoate (11)

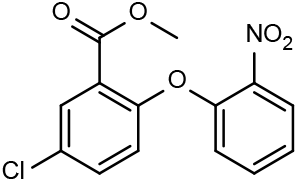

Synthesized according to GP1 from methyl 5-chlorosalicylate (2.80 g, 15.0 mmol), fluoronitrobenzene (1.06 mL, 10.0 mmol) and potassium carbonate (2.07 g, 15 mmol). Purification of the crude product by flash chromatography (*n*-hexane/EtOAc 9:1 to 8:2) yielded the title compound as a colourless solid (3.07 g, 99%). ^1^H NMR (250 MHz, DMSO-*d*_6_) δ 8.07 (dd, *J* = 8.1, 1.7 Hz, 1H) 7.92 (d, *J* = 2.7 Hz, 1H), 7.75 (dd, *J* = 8.8, 2.8 Hz, 1H) 7.68-7.60 (m, 1H), 7.37-7.30 (m, 1H), 7.27 (d, *J* = 8.8 Hz, 1H), 7.00 (d, *J* = 8.4, 1.1 Hz, 1H), 70 (s, 3H).

### Methyl 2-(2-nitrophenoxy)-5-(trifluoromethyl)benzoate (12)

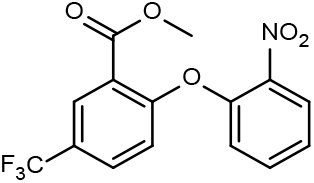

Synthesized according to GP1 from **10** (991 mg, 4.50 mmol), 2-fluoronitrobenzene (317 µL, 3.00 mmol) and potassium carbonate (622 mg, 4.50 mmol). Purification of the crude product by flash chromatography (hexane/EtOAc 9:1 to 8:2) yielded the title compound as a colourless solid (615 mg, 60%). ^1^H NMR (250 MHz, CDCl_3_) δ 8.27 (s, 1H) 8.03 (d, *J* = 8.2 Hz, 1H), 7.75 (d, *J* = 8.6 Hz, 1H), 7.56 (t, *J* = 8.2 Hz, 1H), 7.29 (t, *J* = 8.2 Hz, 1H), 7.08 (d, *J* = 8.6 Hz, 1H), 6.98 (d, *J* = 8.3 Hz, 1H), 3.85 (s, 3H).

### Methyl 2-(2-aminophenoxy)-5-chlorobenzoate (13)

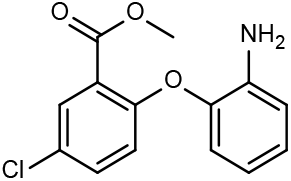

Synthesized according to GP2 from **11** (954 mg, 3.10 mmol) and tin (II) chloride dihydrate (2.80 g, 12.4 mmol). Purification of the crude product by flash chromatography (hexane/EtOAc 9:1 to 8:2) yielded the title compound as a colourless solid (635 mg, 74%). ^1^H NMR (300 MHz, CDCl_3_) δ 7.82 (d, *J* = 2.6 Hz, 1H), 7.33 (dd, *J* = 8.7, 2.6 Hz, 1H), 7.04-6.97 (m, 1H), 6.88-6.80 (m, 3H), 6.71 (td, *J* = 7.7, 1.5 Hz, 1H), 3.89 (s, 3H), 3.64 (br s, 2H).

### Methyl 2-(2-aminophenoxy)-5-(trifluoromethyl)benzoate (14)

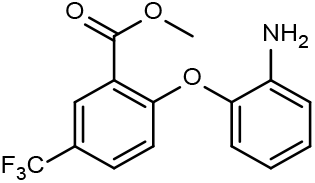

Synthesized according to GP2 from **12** (334 mg, 0.980 mmol) and tin (II) chloride dihydrate (900 mg, 3.91 mmol). Purification of the crude product by flash chromatography (hexane/EtOAc 9:1 to 8:2) yielded the title compound as a colourless solid (171 mg, 56%). ^1^H NMR (250 MHz, CDCl_3_) δ 8.12 (d, *J* = 2.1 Hz, 1H), 7.61 (dd, *J* = 8.7, 2.4 Hz, 1H), 7.10-7.02 (m, 1H), 6.98-6.93 (m, 2H), 6.84 (dd, *J* = 7.9, 1.5 Hz, 1H), 6.79-6.72 (m, 1H), 3.99 (br s, 2H), 3.94 (s, 3H).

### 2-Chlorodibenzo[b,f][1,4]oxazepin-11(10H)-one (15)

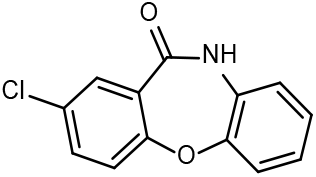

Synthesized according to GP3 from **13** (630 mg, 2.27 mmol) and concentrated sulphuric acid (150 µL, 2.81 mmol). After filtration the title compound was obtained as a colourless solid (508 mg, 90%). ^1^H NMR (250 MHz, DMSO-*d*_6_) δ 10.66 (s, 1H), 7.72 (d, *J* = 2.5 Hz, 1H) 7.67 (dd, *J* = 8.6, 2.8 Hz, 1H), 7.40 (d, *J* = 8.6 Hz, 1H) 7.34 (dt, *J* = 7.1, 1.2 Hz, 1H), 7.21-7.10 (m, 3H).

### 2-(Trifluoromethyl)dibenzo[b,f][1,4]oxazepin-11(10H)-one (16)

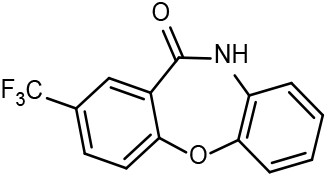

Synthesized according to GP3 from **14** (166 mg, 0.533 mmol) and concentrated sulphuric acid (35.3 µL, 0.663 mmol). After filtration the title compound was obtained as a colourless solid (73 mg, 49%). ^1^H NMR (250 MHz, DMSO-*d*_6_) δ 10.77 (s, 1H), 8.05-7.98 (m, 2H) 7.59 (d, *J* = 8.3 Hz, 1H), 7.39 (dt, *J* = 7.3, 1.2 Hz, 1H), 7.23-7.13 (m, 3H).

### 2-Fluoro-*N*-(5-fluoro-2-hydroxyphenyl)-5-(trifluoromethyl)benzamide (17)

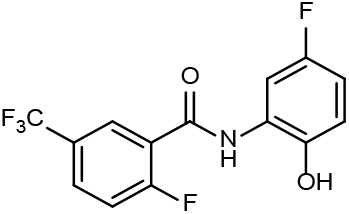

Synthesized according to GP4 from 2-fluoro-5-(trifluoromethyl)benzoic acid (1.50 g, 7.06 mmol), thionylchloride (1.04 mL, 14.1 mmol), 2-amino-4-fluorophenol (925 mg, 7.06 mmol) and triethylamine (1.98 mL, 14.1 mmol). Purification of the crude product by flash chromatography (*n*-hexane/EtOAc 99:1 to 2:3) yielded the title compound as a red solid (1.43 g, 64%). ^1^H NMR (400 MHz, DMSO-*d*_6_) δ 8.17-8.13 (m, 1H), 8.05-7.91 (m, 2H), 7.66-7.59 (m, 1H), 6.93-6.80 (m, 2H).

### 8-Fluoro-2-(trifluoromethyl)dibenzo[b,f][1,4]oxazepin-11(10H)-one (18)

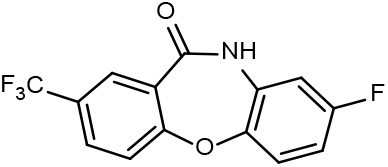

Synthesized according to GP5 from **17** (196 mg, 0.618 mmol), and sodium hydroxide (25 mg, 0.618 mmol). After filtration the title compound was obtained as a light-brown solid (168 mg, 92%). ^1^H NMR (250 MHz, DMSO-*d*_6_) δ 10.85 (s, 1H), 8.04-8.00 (m, 2H), 7.61-7.58 (m, 1H), 7.48-7.40 (m, 1H), 7.06-6.97 (m, 2H).

### 2,11-Dichlorodibenzo[b,f][1,4]oxazepine (19)

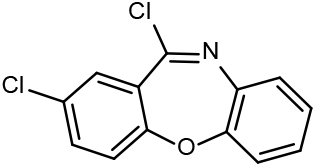

Synthesized according to GP6 from **15** (558 mg, 2.27 mmol), phosphorus oxychloride and *N*,*N*-dimethylaniline (173 µL, 1.36 mmol). After work up the title compound was obtained as brown solid (598 mg, 99%).

### 11-Chloro-2-(trifluoromethyl)dibenzo[b,f][1,4]oxazepine (20)

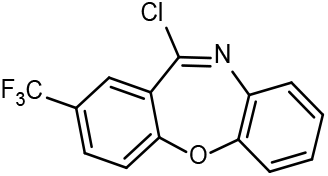

Synthesized according to GP6 from **16** (73 mg, 0.261 mmol), phosphorus oxychloride and *N*,*N*-dimethylaniline (20.3 µL, 0.160 mmol). After work up the title compound was obtained as a brown solid (64 mg, 82%).

### 2,11-Dichlorodibenzo[b,f][1,4]thiazepine (21)

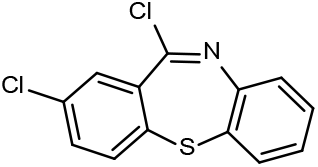

Synthetized according to GP6 from commercially purchased 2-chlorodibenzo[*b*,*f*][1,4]thiazepin-11(10*H*)-one (BLDPharm, Cat. No.: BD223305 97%) (125 mg, 0.478 mmol), phosphorus oxychloride and *N*,*N*-dimethylaniline (36.8 µL, 0.290 mmol). After work up the title compound was obtained as a brown solid (124 mg, 92%).)

### 11-Chloro-8-fluoro-2-(trifluoromethyl)dibenzo[b,f][1,4]oxazepine (22)

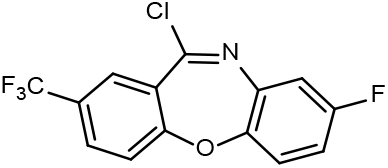

Synthetized according to GP6 from **18** (120 mg, 0.404 mmol), phosphorus oxychloride and *N*,*N*-dimethylaniline (31.0 µL, 0.242 mmol). After work up the title compound was obtained as a light-brown solid. (50.4 mg, 39%).

### 11-(Piperazin-1-yl)-2-(trifluoromethyl)dibenzo[b,f][1,4]oxazepine (23)

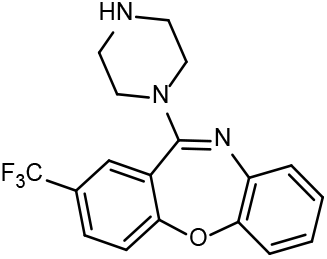

Synthesized according to GP7 from **20** (267 mg, 0.897 mmol) and piperazine (155 mg, 1.79 mmol) in. Purification of the crude product by flash chromatography (DCM/MeOH 95:5 to 9:1) yielded the title compound as a yellow solid (240 mg, 77%). ^1^H NMR (400 MHz, CDCl_3_) δ 8.77 (s, 2H), 8.47 (br s, 1H), 7.73 (d, *J* = 8.0 Hz, 1H), 7.64-7.60 (m, 1H), 7.37 (d, *J* = 8.0 Hz, 1H), 7.18-7.03 (m, 4H), 3.77-3.23 (m, 8H).

### 11-(1,4-Diazepan-1-yl)-2-(trifluoromethyl)dibenzo[b,f][1,4]oxazepine (24)

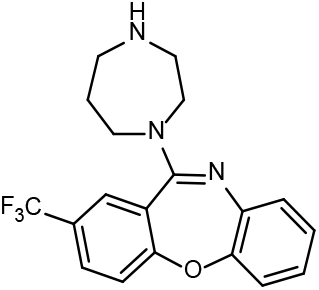

Synthesized according to GP7 from **20** (267 mg, 0.897 mmol) and homopiperazine (183 mg, 1.79 mmol). Purification of the crude product by flash chromatography (DCM/MeOH 95:5 to 9:1) yielded the title compound as a yellow solid (178 mg, 55%). ^1^H NMR (400 MHz, CDCl_3_) δ 7.69 (dd, *J* = 8.6, 2.0, 1H), 7.63 (d, *J* = 1.9 Hz, 1H), 7.36 (d, *J* = 8.5 Hz, 1H), 7.13-7.06 (m, 3H), 7.99-6.95 (m, 1H), 3.98-3.13 (m, 9H), 2.14-1.90 (m, 2H).

### 11-(4-Methylpiperazin-1-yl)-2-(trifluoromethyl)dibenzo[b,f][1,4]oxazepine (1)

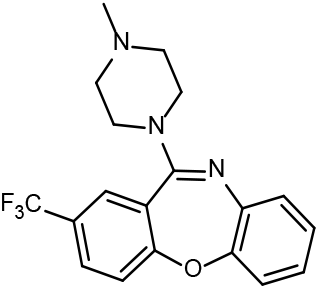

Synthesized according to GP7 from **20** (63 mg, 0.210 mmol) and 1-methylpiperazine (46.6 µL, 0.420 mmol). Purification of the crude product by preparative HPLC yielded the title compound as a brown oil and formate salt (22 mg, 26%). ^1^H NMR (300 MHz, CDCl_3_) δ 8.35 (s, 1H), 7.95 (br s, 1H), 7.70 (ddd, *J* = 8.5, 2.3, 0.5 Hz, 1H), 7.62 (d, *J* = 2.3 Hz, 1H), 7.36 (d, *J* = 8.5 Hz, 1H), 7.18-7.07 (m, 3H), 7.01 (td, *J* = 7.4, 2.0 Hz, 1H), 3.62 (br s, 4H), 2.69 (br s, 4H), 2.44 (s, 3H); ^13^C{^1^H} NMR (75 MHz, CDCl_3_) δ 166.6, 163.2, 158.7, 151.4, 139.9, 129.7, 127.5, 127.2, 127.0, 126.0, 124.9, 124.1, 123.5, 122.1, 120.3, 54.0, 46.7, 45.3; HPLC-purity (254 nm): 96%; MALDI-HRMS: *m/z* calculated for C_19_H_19_F_3_N_3_O[M+H]^+^: 362.1474, found: 362.1488.

### 11-(4-Methyl-1,4-diazepan-1-yl)-2-(trifluoromethyl)dibenzo[b,f][1,4]oxazepine (2)

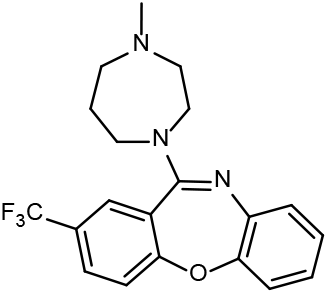

Synthesized according to GP7 from **20** (63 mg, 0.212 mmol) and 1-methylhomopiperazine (49 mg, 0.423 mmol). Purification of the crude product by preparative HPLC yielded the title compound as a brown oil and formate salt (73 mg, 92%). ^1^H NMR (300 MHz, CDCl_3_) δ 9.97 (br s, 1H), 8.51 (s, 1H), 7.68 (dd, *J* = 8.5, 1.9 Hz, 1H), 7.58 (d, *J* = 2.1 Hz, 1H), 7.35 (d, *J* = 8.5 Hz, 1H), 7.12-7.06 (m, 3H), 6.98-6.95 (m, 1H), 4.06-3.45 (m, 4H), 3.08-2.84 (m, 4H), 2.56 (s, 3H), 2.29 (br s, 1H), 2.01 (br s, 1H); ^13^C{^1^H} NMR (75 MHz, CDCl_3_) δ 168.2, 162,8, 158.3, 151.2, 140.4, 129.4, 127.4, 127.0, 126.9, 126.2, 124.2, 124.0, 123.6, 122.2, 120.2, 56.9, 56.3, 49.1, 46.9, 45.4, 26.3; HPLC-purity (254 nm): 97%; MALDI-HRMS: *m/z* calculated for C_20_H_21_F_3_N_3_O[M+H]^+^: 376.1631, found: 376.1639.

### 8-Fluoro-11-(4-methylpiperazin-1-yl)-2-(trifluoromethyl)dibenzo[b,f][1,4]oxazepine (3)

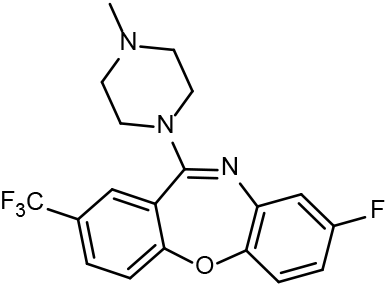

Synthesized according to GP7 from **22** (50 mg, 0.158 mmol) and 1-methylpiperazine (32 mg, 0.317 mmol). Purification of the crude product by preparative HPLC yielded the title compound as a yellow solid (25 mg, 42%). ^1^H NMR (300 MHz, CDCl_3_) δ 7.68 (dd, *J* = 8.5, 1.9 Hz, 1H), 7.58 (d, *J* = 2.1 Hz, 1H), 7.35 (d, *J* = 8.5 Hz, 1H), 7.12-7.06 (m, 3H), 6.98-6.95 (m, 1H), 4.06-3.45 (m, 4H), 3.08-2.84 (m, 4H), 2.56 (s, 3H), 2.29 (br s, 1H), 2.01 (br s, 1H); ^13^C{^1^H} NMR (75 MHz, CDCl_3_) δ 163.2, 161.5, 159.6, 159.4, 147.7, 141.6, 128.4, 127.7, 123.7, 124.1, 122.2, 120.1, 113.4, 110.9, 54.7, 47.3, 46.0; HPLC-purity (254 nm): >99%; MALDI-HRMS: *m/z* calculated for C_19_H_18_F_4_N_3_O[M+H]^+^: 380.1381, found: 380.1377.

### 2-(4-(2-Chlorodibenzo[b,f][1,4]oxazepin-11-yl)piperazin-1-yl)ethan-1-ol (4)

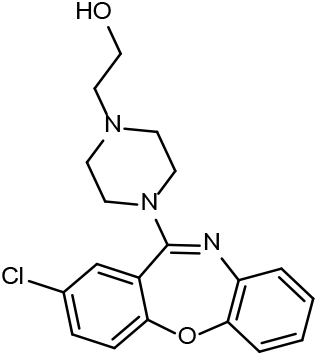

Synthesized according to GP7 from **19** (198 mg, 0.750 mmol) and hydroxyethylpiperazine (195 mg, 1.50 mmol). Purification of the crude product by preparative HPLC yielded the title compound as a brown oil and formate salt (216 mg, 72%). ^1^H NMR (300 MHz, CDCl_3_) δ 8.40 (s, 1H), 7.41 (dd, *J* = 8.6, 2.6 Hz, 1H), 7.31 (d, *J* = 2.6 Hz, 1H), 7.19 (d, *J* = 8.6 Hz, 1H), 7.16-7.07 (m, 3H), 7.05-6.99 (m, 1H), 6.51 (br s, 2H), 3.85-3.81 (m, 2H), 3.74 (br s, 4H), 2.98 (br s, 4H), 2.89-2.85 (m, 2H); ^13^C{^1^H} NMR (75 MHz, CDCl_3_) δ 166.8, 159.3, 158.3, 151.7, 139.6, 133.0, 130.5, 128.8, 127.1, 125.9, 125.1, 124.5, 122.9, 120.2, 59.7, 56.9, 52.2, 45.8; HPLC-purity (254 nm): 99%; MALDI-HRMS: *m/z* calculated for C_19_H_21_ClN_3_O_2_[M+H]^+^: 358.1317, found: 358.1324.

### 2-(2-(4-(2-Chlorodibenzo[b,f][1,4]thiazepin-11-yl)piperazin-1-yl)ethoxy)ethan-1-ol (5)

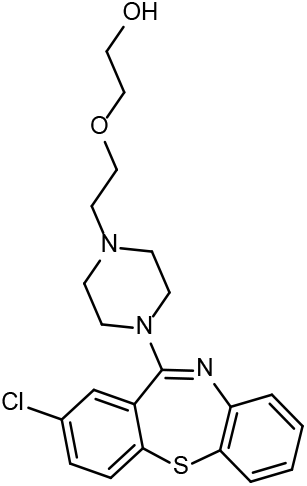

Synthesized according to GP7 from **21** (119 mg, 0.425 mmol) and 1-[2-(2-hydroxyethoxy)ethyl]piperazine (147 µL, 0.850 mmol). Purification of the crude product by preparative HPLC yielded the title compound as a brown oil and formate salt (156 mg, 79%). ^1^H NMR (300 MHz, CDCl_3_) δ 8.37 (s, 1H), 7.46-7.41 (m, 1H), 7.38 (dd, *J* = 7.7, 1.5 Hz, 1H), 7.33-7.27 (m, 2H), 7.29 (br s, 1H), 7.23-7.17 (m, 1H), 7.06 (dd, *J* = 8.0, 1.5 Hz, 1H), 6.93 (td, *J* = 7.3, 1.5 Hz, 1H), 3.88 (br s, 2H), 3.75 (t, *J* = 5.1 Hz, 2H), 3.70 (t, *J* = 4.8 Hz, 2H), 3.58 (t, *J* = 4.8 Hz, 2H), 3.55 (br s, 2H), 3.08-3.00 (m, 2H), 2.90 (t, *J* = 5.3 Hz, 2H), 2.90-2.82 (br s, 2H); ^13^C{^1^H} NMR (75 MHz, CDCl_3_) δ 166.7, 158.9, 148.2, 138.1, 134.9, 134.8, 133.4, 132.9, 131.2, 129.4, 128.6, 127.3, 125.3, 123.6, 72.6, 66.1, 61.4, 57.2, 52.0, 45.1; HPLC-purity (254 nm): 95%; MALDI-HRMS: *m/z* calculated for C_21_H_25_ClN_3_O_2_S[M+H]^+^: 418.1351, found: 418.1350.

### 2-(2-(4-(2-Chlorodibenzo[b,f][1,4]oxazepin-11-yl)piperazin-1-yl)ethoxy)acetic acid (6)

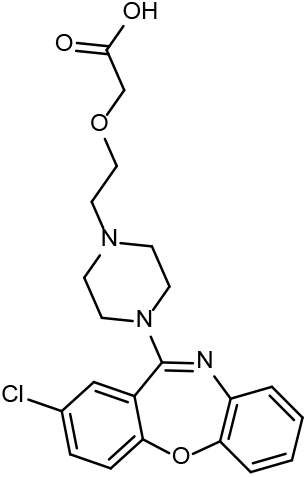

Synthesized according to GP8 from commercially purchased amoxapine (TCI, Product Number: A2499) (75 mg, 0.239 mmol), 2-(2-chloroethoxy)acetic acid (66 mg, 0.478 mmol) and triethylamine (0.34 mL, 2.39 mmol). Purification of the crude product by preparative HPLC yielded the title compound as a brown oil and formate salt (47 mg, 42%). ^1^H NMR (300 MHz, CDCl_3_) δ 8.24 (br s, 1H), 7.41 (dd, J = 8.7, 2.4 Hz, 1H), 7.34 (d, J = 2.4 Hz, 1H), 7.18 (d, J = 8.7 Hz, 1H), 7.15-6.98 (m, 4H), 5.91 (br s, 2H), 4.23 (br s, 2H), 4.44-3.94 (m, 6H), 3.87-3.72 (m, 6H); ^13^C{^1^H} NMR (75 MHz, CDCl_3_) δ 168.8, 164.7, 159.3, 159.0, 151.6, 139.1, 133.4, 130.8, 128.7, 127.2, 126.0, 125.7, 123.9, 123.0, 120.3, 61.4, 59.5, 55.8, 41.4; HPLC-purity (254 nm): 99%; MALDI-HRMS: *m/z* calculated for C_21_H_23_ClN_3_O_4_ [M+H]^+^: 416.1372, found: 416.1375.

### 2-(2-(4-(2-(Trifluoromethyl)dibenzo[b,f][1,4]oxazepin-11-yl)piperazin-1-yl)ethoxy)acetic acid (7)

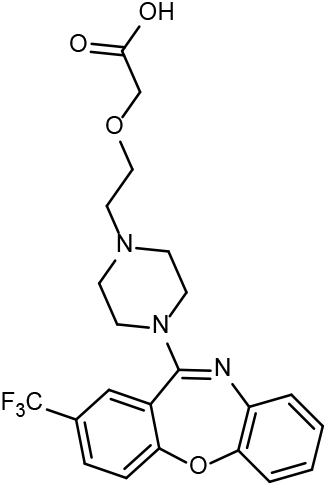

Synthesized according to GP8 from **23** (75 mg, 0.102 mmol), 2-(2-chloroethoxy)acetic acid (28 mg, 0.205 mmol) and triethylamine (0.14 mL, 1.02 mmol). Purification of the crude product by flash chromatography (DCM/MeOH 95:5 to 9:1) yielded the title compound as a colourless solid (28 mg, 61%). ^1^H NMR (400 MHz, CDCl_3_) δ 7.66 (dd, J = 8.5, 2.0 Hz, 1H), 7.56-7.56 (m, 1H), 7.31 (d, J = 8.5 Hz, 1H), 7.12-7.03 (m, 3H), 6.97 (td, J = 7.6, 1.8 Hz, 1H), 4.21 (s, 2H), 3.73-3.43 (m, 12H); ^13^C{^1^H} NMR (101 MHz, CDCl_3_) δ 168.7, 163.4, 158.8, 151.5, 139.7, 128.5, 127.8, 127.4, 126.3, 125.4, 124.1, 123.6, 122.5, 120.5, 73.8, 69.7, 61.8, 44.3, 41.7; HPLC-purity (254 nm): 98%; MALDI-HRMS: *m/z* calculated for C_22_H_23_F_3_N_3_O_4_ [M+H]^+^: 450.1635, found: 450.1637.

### 2-(2-(4-(2-(Trifluoromethyl)dibenzo[b,f][1,4]oxazepin-11-yl)-1,4-diazepan-1-yl)ethoxy)acetic acid (8)

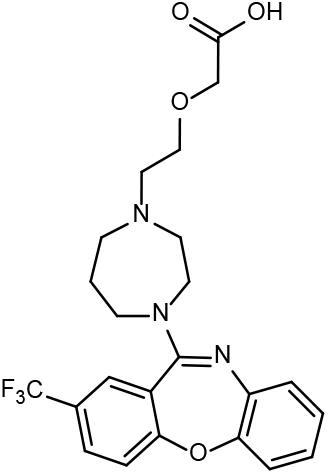

Synthesized according to GP8 from **24** (40 mg, 0.111 mmol), 2-(2-chloroethoxy)acetic acid (31 mg, 0.221 mmol) and triethylamine (0.16 mL, 1.11 mmol). Purification of the crude product by flash chromatography (DCM/MeOH 95:5 to 9:1) yielded the title compound as a colourless solid (20 mg, 39%). ^1^H NMR (400 MHz, CDCl_3_) δ 7.95-7.91 (m, 1H), 7.80-7.77 (m, 1H), 7.59 (t, J = 8.8 Hz, 1H), 7.22-7.18 (m, 1H), 7.11-6.93 (m, 3H), 4.63 (d, J = 4.8 Hz, 1H), 4.18-4.09 (m, 3H), 3.77-3.36 (m, 10H), 2.08-1.46 (m, 2H); ^13^C{^1^H} NMR (101 MHz, CDCl_3_) δ 168.6, 162.4, 157.6, 150.7, 140.3, 129.9, 126.6, 126.3, 126.0, 126.0, 123.7, 123.6, 123.7, 122.4, 120.2, 72.5, 69.1, 68.9, 68.2, 67.6, 63.2, 60.0, 44.0; HPLC-purity (254 nm): 95%; MALDI-HRMS: *m/z* calculated for C_23_H_25_F_3_N_3_O_4_ [M+H]^+^: 464.1792, found: 464.1788.

